# Life cycle complexity shapes parasite sharing amongst migratory and resident hosts

**DOI:** 10.64898/2026.01.14.699309

**Authors:** Sarah Nichols, Andrea Estandía, Catherine M Young, Jaslyn Allnutt, Jonathan T Coleman, Judith Little, Gregory Little, Sonya M Clegg, Beth Okamura

## Abstract

Migratory animals are potentially important parasite dispersers. However, understanding the impact of host migration on parasite distributions is challenging because variation in parasite life histories can mediate dispersal potential. Furthermore, studies of parasite dispersal via host migration often compare parasites in different species of migrant and resident hosts, despite differences in host phylogeny, ecology, and exposure history, which can complicate ascertaining the impact of migration. The Tasmanian population of silvereye (*Zosterops lateralis lateralis*) exhibits partial migration. Some members of the population overwinter on the Australian mainland, where they co-occur with residents of another silvereye subspecies (*Z. l. cornwalli*). Utilising targeted PCR, metabarcoding and high-throughput sequencing, we characterise a range of parasite genera from blood and faecal samples in Tasmanian resident, migrant and mainland resident birds. Greater parasite richness was documented in migrant individuals, possibly reflecting exposure to varied parasite faunas at stopovers or during overwintering. The increased richness was primarily driven by parasites with simple life cycles. Reduced sharing of parasites with complex life cycles amongst migratory and resident host populations may be explained by ecological constraints in establishing transmission, such as vector or secondary host availability. Our study exemplifies the utility of molecular methods for characterising multi-parasite systems and makes use of a single host species to provide insights on patterns of parasite distributions that result from migratory movements. Knowing how parasite life histories constrain or promote dispersal via regular host migration will be essential for predicting future patterns of pathogen dispersal under climate change and shifting migratory regimes.

## Introduction

Migratory animals embarking on seasonal long-distance movements typically encounter a variety of habitats and organisms, exposing them to a wide array of pathogens and parasites. Hosts may consequently lose, swap or gain parasites as a result of their seasonal migratory movements (1; 2). Such migration-related exposure to parasites can profoundly impact the dynamics of infectious disease (3). Yet, the impact of migration on parasite dispersal and distribution remains difficult to predict, particularly because processes that cause migratory hosts to gain and lose parasites may be happening concurrently, and because similar patterns of parasite distributions can arise via different mechanisms (4). With global climate change altering the timing, routes, and scale of animal migrations (5; 6; 7; 8), elucidating the impact of host migration on parasite distribution has become increasingly urgent (9; 10).

Several hypotheses have been established regarding dynamics of parasites in migratory hosts. Hosts may gain a diverse parasite community via exposure to multiple parasite assemblages during stopovers or upon arrival, i.e. “migratory exposure” (11; 12). Parasites may also alter migratory host behaviours by imposing energetic costs which lead to trade-offs among functions (13). For example, physiological trade-offs with the immune system (14), may result in less variation at the immune loci (15), and consequently higher parasite loads in migratory animals (3). Contrastingly, migrants have also been hypothesised to have evolved varied immune loci as a response to frequent parasite exposure (16), resulting in them having a less diverse parasite community. Alternatively, the energetic cost of migration may lead to behavioural tradeoffs. Thus, the “migratory escape” hypothesis proposes that periodic migration enables hosts to avoid the accumulation of parasites (17). The detrimental impact of infection on host fitness may also lead to loss of parasites from host populations via “migratory culling” (18; 19). Such evolutionary responses from hosts may, in turn, lead to changes in parasite phenology (20), transmission, or survival strategies (21). Disentangling the effects of these non-mutually exclusive hypotheses is challenging, particularly as they may operate in tandem with different parasite genera imposing opposing effects within the same host (22).

Empirical studies that compare parasite diversity in migratory and resident birds have produced varied results. Greater parasite richness in migratory compared to resident host species have been documented for nematodes (11; 12); whereas for haemosporidians, the causative agents of avian malaria, results are often contradictory. Some studies have found higher richness in migratory bird species (23) while others have found no detectable difference between resident and migrant species (24).

Quantifying parasite diversity in migratory hosts is important for understanding the potential for migration-facilitated parasite sharing. In terms of sharing at the host species community level, studies have yielded mixed results. Perrin *et al*., (25) compared 697 bird species and found that migratory species shared many *Haemoproteus* and *Plasmodium* lineages with resident species encountered along the African-Eurasian and Americas flyways, implying that migrants may disperse and/or acquire parasites during their journeys. Another study, based on 896 bird species sampled across South America revealed that communities with a higher proportion of migrant birds supported higher haemosporidian lineage richness (26). In contrast, other studies have found that various species of co-occurring migratory and resident Passeriformes harbour largely distinct haemosporidian communities at stopover and overwintering sites in South and North America (27; 28; 29). Such results may reflect differences in vector availability or competency across the migrant hosts’ annual cycle. For example, Galen *et al*., (29) examined haemosporidian parasites in 63 species of birds at a stopover site in Pennsylvania (USA) and found that migratory birds shared a core set of haemosporidians with residents. However, migratory bird species also carried several unique haemosporidian lineages associated with different biogeographic regions that were characterised by different vector communities. This suggests that vector ecology may preclude transmission for some haemosporidian parasites. Such results highlight that dispersal of parasites with complex life cycles may be constrained by the availability of secondary hosts or vectors. Therefore, it is important to assess parasites with differing life cycles when considering the impact of migration on parasite sharing.

To date, studies that attempt to quantify parasite diversity and subsequent sharing between resident and migrant hosts often rely on species-level comparisons. Such comparisons do not necessarily account for differences in host phylogeny, ecology, and exposure history, which may confound efforts to resolve the effects of migration on parasite communities (30). Partial migrant systems, in which individual-level variation in migratory behaviour occurs, provide valuable opportunities to understand parasite transmission dynamics during migration in a single host species. Although partial migration is relatively common in birds (31; 32), distinguishing migrants from non-migrants without individual tracking can be challenging. However, this issue can be avoided by strategically sampling during the migratory period, when migrant and resident birds self-sort. Furthermore, systems in which migrating individuals encounter another morphologically distinct sub-species during the migratory period enable further resident and migrant comparisons to be made. Such systems offer a tractable framework for studying parasite sharing between resident and migratory populations.

The Tasmanian population of silvereye (*Zosterops lateralis lateralis*), a passerine bird of Australia and the southwest Pacific, undergoes partial migration (33). During the winter, some individuals remain on the Tasmanian breeding grounds while others cross the Bass Strait to overwinter on the Australian mainland, where they co-occur with year-round residents of another morphologically distinct subspecies of silvereye (*Zosterops lateralis cornwalli*). We take advantage of this system of resident and partial migrant silvereye sub-species to understand how migration shapes parasite communities. By utilising high-throughput sequencing (HTS) methods we characterise a broad range of parasite genera with varied life-histories detected in host faeces and blood. This enables us to directly address whether differences in parasite richness between resident and migrant birds depend on life cycle complexity, and if some parasites are more broadly shared between host groups than others.

## Materials and Methods

### Sample collection

We conducted sampling from May to July 2022 when the Tasmanian population of silvereye (*Z. l. lateralis*) undergoes partial migration (33). During May and June, samples from overwintering migratory individuals were collected in Queensland (QLD) and New South Wales (NSW) on the Australian mainland. We also collected samples from individuals of the year-round resident subspecies, *Z. l. cornwalli*, which were present in the same area as migrants. Individuals of the two subspecies were discriminated by plumage differences (34). Samples from individual Tasmanian residents were collected during June 2022. Figure. 1A illustrates the sampling locations and partial migrant route.

**Fig. 1.**
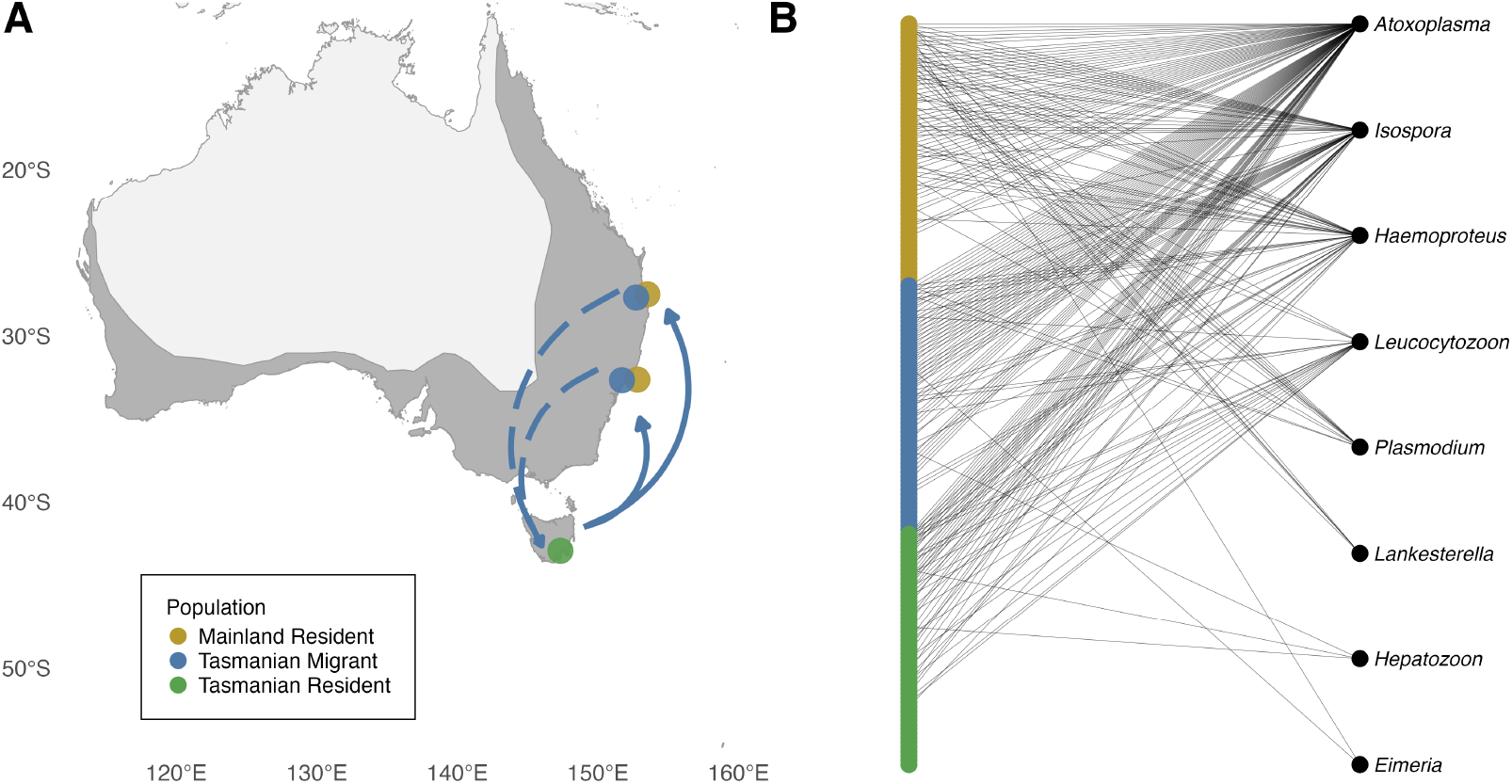
A) Map showing silvereye distribution in Australia (in dark grey) and sampling (indicated by circles) for Tasmanian residents (green, n = 53), Tasmanian migrants (blue, total n = 56, comprising New South Wales [NSW] = 41 and Queensland [QLD] = 15) and mainland residents (yellow, total n = 59, comprising NSW = 43 and QLD = 16), the blue arrows indicate the north-south migrant movement, Tasmanian migrants were sampled after they arrived on the mainland (direction of travel indicated by solid blue line), before their presumed return to Tasmania (dashed line). B) Corresponding bipartite network constructed for 8 parasite genera and 168 silvereye individuals from the three silvereye populations.

Faecal samples were collected according to a protocol adapted from Knutie and Gotanda (35). Birds were captured using a mist net and subsequently placed in a ventilated, flat-bottomed paper bag lined with two sterile weigh-boats covered by wire mesh that acted as a perch (Figure S1). Individual birds were left in the bag for approximately three minutes to defecate. Faecal samples were transferred to 1.5 ml microfuge tubes containing 1 ml 100% ethanol. A blood sample was taken by pricking the brachial vein with a sterile needle and collecting the resulting blood droplet in a glass capillary tube. Capillary tubes were snapped to size then placed in 1.5 ml microfuge tubes with Queen’s lysis buffer (0.01 M Tris, 0.01 M NaCL, 0.01 M EDTA and 0.03 M n-lauroylsarcosine (10% w/v), pH 7.5, modified from Seutin *et al*., (36). All samples were refrigerated until they could be frozen at −20°C. Birds were fitted with an individually numbered band provided by the Australian Bird and Bat Banding Scheme. A suite of morphometrics were measured with dial calipers ±0.1 mm (tarsus length as well as bill length, depth and width) and a flat rule ±1 mm (wing length and tail length). Weight was measured with a Pesola scale. Birds were then released.

### DNA extractions

Faecal DNA extractions were carried out using the Qiagen DNeasy PowerSoil Pro Kit (Qiagen, Germany) according to the manufacturer’s instructions. Blood extractions were conducted using a modified phenolchloroform protocol (37). Samples were incubated overnight at 55°C with rotation, adding approximately 100 µl of blood to 250 µl DIGSOL extraction buffer (0.02 M EDTA, 0.05 M Tris-HCl [pH 8.0], 0.4 M NaCl, 0.5% sodium dodecyl sulphate [SDS]) and 10 µl Proteinase K (20 mg/ mL). Digested samples were added to 250 µl phenol:chloroform:isoamyl alcohol (25:24:1) and rotated for 10 minutes followed by centrifugation at 10,000 rpm for 10 minutes. The aqueous phase was transferred to a new microfuge tube; rotation and centrifugation were repeated with the addition of 250 µl phenol:chloroform:isoamyl alcohol (25:24:1). This step was then repeated with the addition of 250 µl chloroform:isolamyl alcohol (24:1); and 2 volumes of cold 100% ethanol, 1 volume of 2.5 M ammonium acetate and 1 µl glycogen to precipitate the DNA. Samples were stored at −20°C for at least 12 hours before further centrifugation for 10 min at 15,000 rpm at 4°C, the precipitate was washed with 500 µl cold 70% ethanol. The process of centrifugation and washing was done twice more. The precipitated DNA was left to dry at room temperature before resuspension in 25 µl of TE (TrisEDTA) buffer (0.01 M Tris-HCL (pH 8.0), 0.0001 M EDTA). Each batch of 24 samples included one negative extraction control.

### Sexing

Silvereyes are sexually monomorphic (38), therefore individuals were sexed genetically. We used two sets of markers: P2D-P8 (39) and Z002B (40). PCR amplification was conducted in a total volume of 2 µl, consisting of 1 µl QIAGEN Multiplex PCR mix (5X, QIAGEN Inc.) and 1 µl of primer mix, containing 5 µM of each primer and ddH2O and 15 ng of DNA, obtained by allowing approx. 1 µl of resuspended DNA extract to evaporate. The PCR conditions involved an initial denaturation stage at 95°C for 15 min, followed by amplification for 35 cycles (94°C for 30 s, 48°C [P2D-P8] or 56°C [Z002B] for 90 s and 72°C for 60 s, and a final extension for 30 min at 60°C. PCR products were diluted in ddH2O to a ratio of 1:2500-1:5000 and analysed on an ABI 3730 48-capillary DNA Analyser using formamide and GeneScan™-500 ROX size-standard (Applied Biosystems, Warrington, UK). Alleles were scored using GENEMAPPER v 5 software (Applied Biosystems, California, USA).

### 18S anti-metazoan metabarcoding

A set of 18S anti-metazoan primers, 3NDF (41) and the “universal non-metazoan” reverse primer, 18s-EUK1134-R (42), were used to amplify the V4 region of the 18S SSU rDNA. We chose these primers because the 18S region is highly conserved across taxa, but shows substantial variation among taxonomic groups (43; 44; 45). Furthermore, this region is frequently used to characterise parasites (46), providing comparatively good representation in reference databases compared to other regions. The primers were designed to avoid amplifying metazoans as otherwise the resultant sequencing reads would likely be dominated by the host. The primers were modified to include annealing sites for indexing. The primer sequences were as follows:

Forward primer:

5’-ACACTCTTTCCCTACACGACGCTCTTCCGATCTNNNNNGCAAGTCTGGTGCCAG-3’

Reverse primer:

5’-GTGACTGGAGTTCAGACGTGTGCTCTTCCGATCT CTTTAARTTTCASYCTTGCG-3’

PCR1 reactions had a final volume of 20 µl and included Qiagen 5X Multiplex PCR Master Mix (Qiagen, Germany), 2 µl of primer at 5 µM (final conc. 0.5M), 1 µl DNA (ca. 10 ng/µl) and sterile ddH2O. The thermal cycling conditions were as follows: 95°C for 15 min, 35 cycles of 95°C for 30 sec, 53°C for 45 sec and 72°C for 1 min with a final extension at 72°C for 10 min. Two negative controls were included for every 30 amplifications: a PCR control (using 1 µl ddH2O instead of DNA) and an extraction control (using the negative controls generated during DNA extractions). PCR reactions produced a 540bp product.

### Library preparation and sequencing

Successfully amplified samples were cleaned with Promega ProNex Magnetic Beads using a ratio of 1.15X (v/v) beads to product, according to the manufacturer’s instructions (Promega, Inc., United States). Each sample was dual indexed using a PCR reaction that included 5X Meridian Bioscience MyTaq HS Mix, Fi5 and Fi7 primers (at 10 µM), sterile ddH2O and product from PCR1 in a total volume of 20 µl. The PCR conditions were 95°C for 15 min; 8 cycles of 98°C for 10 sec, 65°C for 30 sec and 72°C for 30 sec and a final extension at 72°C for 5 min. We used the Agilent TapeStation (Agilent Technologies Inc., United States) to confirm the index primers had successfully ligated and subsequently conducted a second clean-up using a 1X (v/v) ratio of Promega ProNex Magnetic Beads. The concentration of cleaned product (ng/µl) was quantified using the Promega QuantiFluor and pools were created by combining equal ratios of amplicons from each sample. Each resultant pool was further quantified by qPCR. A 100-1000- and 10000-fold dilution was prepared for each pool with sample dilution buffer (10mM Tris, pH 8.0 and 0.05% Tween 20X) and SYBR FAST mix and primers. The thermal cycling conditions were 5 min at 95°C, 35 cycles of 30 sec at 95°C and 45 sec at 60°C. Pools were further amalgamated and quantified by qPCR until a single pool remained. A High Sensitivity D1000 ScreenTape was used to ascertain the size and quantity of the final product using an Agilent TapeStation.

The final pool was sequenced on a PacBio Sequel II at the NERC Environmental Omics Facility, University of Liverpool, UK. Library loading was achieved by diffusion, with all the samples loaded onto one SMRT cell 8 M, using the Sequel II Primer V3.1.1.5, Sequel®II Binding Kit V3.1. PacBio CCS reads (mean passes = 20, median read quality = Q37) were demultiplexed and Illumina primers were removed using the cutadapt software (47).

### Calling ASVs and taxonomic assignment

The DADA2 package (48) was used to clean and call Amplicon Sequence Variants (ASVs) from the data. Reads with a minimum quality score of Q30 and those within the minimum (475 bp) and maximum (675 bp) lengths were retained. Following filtering, reads were dereplicated. Dereplicating reduces computation time by amalgamating unique sequences, while retaining information about their abundance in the dataset. The samples were subsequently denoised using the learned error rates to remove likely sequencing errors. Chimeras were also removed.

Taxonomic identification of metabarcoding data is often based on sequence similarity. However, sequence similarity ignores evolutionary context (49) and is known to give rise to erroneous identifications (50), particularly when there are limited reference sequences available (51). For these reasons, we utilised phylogenetic placement to taxonomically assign ASVs. Phylogenetic placement requires a reference tree which represents sequences spanning the genetic diversity likely to be present in a sample (49). We used general 18S anti-metazoan primers; therefore, we needed to include a range of eukaryotic sequences. To do this, we utilised the 211 sequences curated by Rapp and Wolf (52) spanning the eukaryotic tree of life, supplemented with additional parasitic sequences that we have previously identified in silvereyes (53). A full list of accession numbers used to build the tree is available in Table S1. We trimmed the reference sequences to encompass the region amplified by our primers using cutadapt (47) and subsequently removed any sequences that exceeded 1000 bp as these were assumed to be erroneous (expected amplicon size is 600 bp). Next, we created a multiple sequence alignment (MSA) using MAFFT (54) and trimmed aligned sequences using trimAL (55), removing any alignsment columns where more than 50% of sequences had gaps. We used ModelTest-NG to select a substitution model (GTR + G4) (56) and inferred a phylogenetic tree using RAxML-NG (referred to as a maximum likelihood (ML) tree) (57). The alignment was reduced for identical sequences and a ML tree search was conducted with 25 parsimony and 25 random starting trees. Felsenstein bootstrap proportion (FBP) was calculated from 1000 bootstrap replicates to assess node support (58). We did not root the tree as this is not required for phylogenetic placement (49). Next, we aligned ASVs to the reference MSA using PaPaRa (59), placed them onto the reference tree using EPA-ng (51) and finally, converted placements to taxonomic assignments with gappa (60). Taxonomic assignments with a Likelihood Weight Ratio (LWR) of 0.7 were retained for further analysis.

After taxonomic assignment, we used the microDecon package to identify and remove potential contaminants, using the 26 sequenced negative controls generated during the PCR and extraction steps as references (61). To account for variation in sequencing depth amongst samples, we standardised read depth by rarefying each sample to a minimum sample coverage of 0.9, following the approach recommended by Chiu *et al*., (62).

We searched the available literature to compile life history information for each of the identified parasite genera (Table S2). Specifically, we collated information on previously recorded primary hosts according to the following categories: bird, other vertebrate, invertebrate or plant. We excluded parasites whose life cycle has not been elucidated in vertebrates. Similarly, we recorded information about the parasite life cycle, categorising parasites into simple (requiring one host) or complex (requiring multiple hosts).

### Haemosporidian-specific multiplex PCR

Despite the availability of reference sequences in public database, 18S metabarcoding was unable to detect haemosporidians (*Haemoproteus, Plasmodium* and *Leucocytozoon*). This could be because amplification of mixed template PCRs is biased against AT-rich sequences (63) and haemosporidian genomes are known to be AT-rich (64). Therefore, we used a multiplex PCR assay which utilises equimolar concentrations of three sets of primers, targeting different genomic regions of each parasite, these are: PMF, PMR, HMF, HMR, LMF, and LMR (65). Each reaction had a final volume of 10 µl and included Qiagen 5X Multiplex PCR Master Mix (Qiagen, Hilden, Germany), 1 µl of each primer at 10 µM (final conc. 0.5M), 1 µl DNA (ca. 10 ng/µl) and sterile ddH2O. The thermal cycling conditions were as follows: 95°C for 15 min; followed by 35 cycles of denaturation at 94°C for 30 s, annealing at 59°C for 90 s, and extension at 72°C for 30 s; with a final extension at 72°C for 10 min. We included one negative (ddH2O) and one positive control for every 16 samples. For the positive control, we alternated between using DNA extracts taken from birds with a confirmed *Haemoproteus, Leucocytozoon*, and a mixed *Haemoproteus* and *Plasmodium* infection. Amplified products were resolved using electrophoresis for 1 h at 90 V on a 2% agarose gel containing GelRed™gel stain (Biotium, Inc., Hayward, CA, USA). The product size was used to infer which genera were present in each sample (*Haemoproteus*: 533 bp, *Plasmodium*: 378 bp, *Leucocytozoon*: 218 bp). For some analyses, we used presence-absence data for the parasite genera detected per individual by our metabarcoding approach and combined it with the resulting information on haemosporidian incidence.

### Statistical analysis

#### ASV-based analyses

All statistical analyses were conducted in R version 4.4.2 (66). To test whether migrant birds carry a greater richness of parasites than their resident conspecifics, we estimated richness for each individual using the standardised ASV data. We estimated observed richness (number of ASVs detected) and asymptotic Chao1 richness per sample type (faecal, blood and combined) using the iNEXT package (67). To quantify whether the migrant population had compositionally distinct parasite communities compared to Tasmanian and mainland resident populations, we calculated Local Contribution to Beta Diversity (LCBD). LCBD can be used to quantify proportional contributions to overall beta diversity within the dataset (68). To compute LCBD, we generated a Jaccard dissimilarity matrix from the standardised ASVs which we had converted to presence-absence data, and calculated LCBD using the Adespatial package (69). Differences in parasite richness (Chao1 and observed richness) as well as community distinctiveness (LCBD) among the populations were assessed using Kruskal-Wallis tests and subsequent post hoc Dunn’s tests. We used Holm-adjusted p-values to account for multiple comparisons.

Next, we tested for differences in the parasite communities sampled in mainland resident and Tasmanian migrant populations in QLD and NSW. Southern to central-eastern QLD is considered the limit for silvereye migration (70; 71; 72). Therefore, birds sampled in QLD are more likely to have reached their overwintering sites than those sampled in NSW. Consequently, a difference in parasite community richness between the two states (QLD and NSW) could help to elucidate when parasites are gained during the migratory cycle. We calculated Chao1 and observed richness for mainland residents and Tasmanian migrants sampled in each state. Then, we fit two generalised linear models (GLM) with a negative binomial error distribution using Chao1 and observed richness as the response variables. We included the following factors as possible predictors: sex, date of sampling, population and sampling state. We included an interaction term between population and sampling state to test whether parasite richness in the Tasmanian migrants differed between the states.

#### Network-based analysis

We built a bipartite network using the presence-absence data derived from metabarcoding and the haemosporidian-specific PCR. The bipartite network enables us to explore whether parasite life cycle complexity influenced patterns of sharing among populations. We split the parasite genera into two categories depending on their life cycle complexity. To test whether parasite life cycles had an impact on richness of parasites infecting the three host populations, we used the networks to calculate scaled degrees per life cycle complexity (i.e. the number of links from each bird to parasite genera with simple or complex life cycles, standardised according to the number of total parasite genera in the network). We fitted a GLM using scaled degrees as the response variable, implemented with a binomial error distribution. The following factors were included as possible predictors: sex, date of sampling, population. We included an interaction term between parasite life cycle complexity and population to test whether parasites with simple life cycles more readily infected migrants. To account for repeated measures per host individual (for simple and complex parasite genera), we report cluster-robust standard errors. Host individual could not be included as a random effect because effective variation was 0, indicating no detectable within-individual variation.

We included sex and date of sampling in all our models. This is because previous work has shown sex-specific variation in parasite prevalence (73; 74; 75; 76) and migratory behaviour (77; 78; 79). We included date of sampling to account for potential seasonal variation in parasite dynamics, as we sampled birds from May to July, during the Austral autumn and winter. Parasite prevalence, load and diversity have been observed to vary in migrant hosts over time (80; 81; 82), with late arrivals reported as having both higher (83; 84) and lower parasite burdens in different systems.

Body condition has also been shown to vary with parasite infection status (86; 87) as well as over the course of migration (88) and with migratory distance (89). Therefore, it is of interest to test whether host body condition impacted individual parasite communities. However, we were unable to include body condition in our GLMs due to missing weight data for 29 individuals. Therefore, to address whether body condition could have impacted our results we conducted a separate GLM with binomial error distribution using scaled degrees (extracted from the bipartite network, representing proportion of total parasite genera detected per host individual) as the response variable. We included body condition, measured by extracting the residuals from an Ordinary Least Squares regression of body mass (g) by tarsus length (cm) (90); sex; population; and date of sampling as possible predictors.

To avoid inclusion of collinear predictor variables in the models, variance inflation factors (VIFs) were computed using the performance package to quantify multicollinearity (91). All predictor variable scores were below three, indicating acceptable levels of collinearity for inclusion in the models (Table S3). Model fit was assessed using the performance package (Figure S2, S3, S4 and S5).

We performed model selection and averaging using the MuMIn package (92). All candidate models were compared based on Akaike’s Information Criterion corrected for small sample sizes (AICc), and model selection tables are presented in Table S4, S5, S6 and S7. Models within ΔAICc <2 of the top-ranked model were considered equally plausible and we generated model-averaged predictions across this top model set.

To interpret the output of our model selection and averaging approach, we consider the weighted importance (WIP) value, which reflects how frequently a predictor appears in the top-ranked models, weighted by the Akaike weights of those models. A WIP close to 1.0 indicates that the variable is consistently included in high-support models, and likely helps to improve model fit. The 95% CI indicates the range of possible values for the coefficient estimate. When the 95% CI include zero, there is insufficient information in the data to rule out a positive or negative effect, and the biological interpretation for that predictor is therefore less clear.

We tested whether certain parasite genera tended to infect a broader range of host individuals than would be expected by chance. We used the bipartite network to calculate proportional similarity per parasite genera (i.e. specialisation within the network, measured as similarity between host use and availability). We compared the z-scores for proportional similarity to the range of z-scores generated in a fixed-fixed null model. In a fixed-fixed null model, the number of parasite infections and host individuals remains the same, but the incidence of infection is randomly allocated in the network.

## Results

We identified parasites in 168 individuals comprising 53 Tasmanian residents, 56 Tasmanian-origin migrants and 59 mainland residents (Figure. 1B). Eight parasite genera with life cycles characterised in vertebrate hosts were recovered (Table S2): five from metabarcoding analysis and three haemosporidian genera from targeted PCR. Three parasite genera are considered to have simple life cycles (*Eimeria* (93), *Isospora* (94; 95; 96), and *Atoxoplasma* (97)) and five to have complex life cycles (*Haemoproteus, Plasmodium, Leucocytozoon* (98), *Lankesterella* (99), and *Hepatozoon* (100)).

Parasite richness varied among the populations (Figure. 2). There were significant differences among populations when comparing ASV sequences extracted from faecal samples for Chao1 (H_df =2_ = 6.91, p = 0.032) and observed richness (H_df =2_ = 12.32, p = 0.002) as well as when combining both sample types for Chao1 (H_df =2_ = 7.69, p = 0.021) and observed richness (H_df =2_ = 6.05, p = 0.049). However, no significant difference was observed for blood parasites for Chao1 (H_df =2_ = 0.43, p = 0.807) or observed richness (H_df = 2_ = 1.82, p = 0.402). There was also a significant difference in LCBD (i.e. proportional contributions to overall beta diversity) for each population for faecal (H_df =2_ = 8.00, p = 0.018) and the combined samples (H_df =2_ = 5.75, p = 0.056) but not blood (H_df =2_ = 0.22, p = 0.895). The Tasmanian migrant population had higher richness than the Tasmanian residents for Chao1 faecal and combined samples as well as observed richness for faecal samples. The mainland residents also had higher richness than the Tasmanian residents for faecal and combined samples according to observed richness and when samples were combined according to Chao1 (Figure. 2). Table S8 summarises the pairwise comparisons generated with post-hoc Dunn tests.

**Fig. 2.**
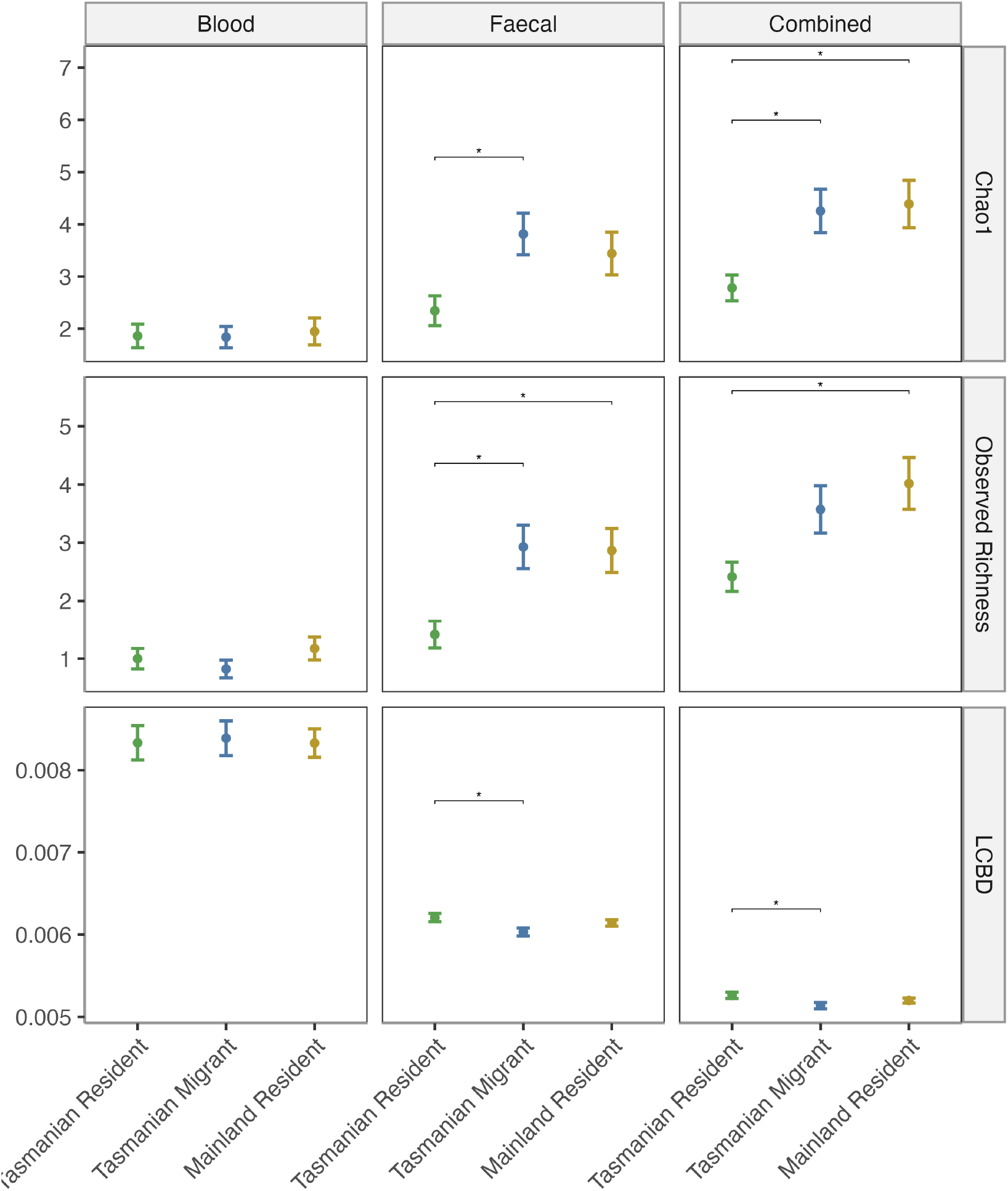
Mean estimated Chao1, Observed Richness and Local Contribution to Beta Diversity (LCBD), (± SE) for each population (Tasmanian residents, Tasmanian migrants and mainland residents) according to standardised ASV data extracted from blood and faecal samples and both types of samples combined. Significant population comparisons indicated by square brackets and asterisks, calculated according to post-hoc Dunn’s test with Holm correction (P < 0.05).

When testing the difference in parasite richness between mainland resident and Tasmanian migrants for each state, we identified three and four models within AICc <2 for each model utilising Chao1 and observed richness as the response variable, respectively (Table S4 and S5). Both models found that state was an important predictor of parasite richness attributing state a WIP of 1 (Table 1). In both cases, richness was higher in the NSW population (Figure. 3). However, neither top model set retained the interaction term between state and population, implying that there was no difference in richness detected in the Tasmanian migrants in each mainland state. Both sets of models also supported the inclusion of sex as a predictor, with WIP of 0.20 (Chao1) and 0.37 (observed richness) although there was some uncertainty about the scale and direction of the effect.

**Table 1.** Model-averaged estimates for General Linear Model predicting Chao1 and observed richness for Tasmanian migrants (TM) and mainland residents (MR) sampled in New South Wales (NSW) and Queensland (QLD). Coefficient estimates (β), standard errors (SE), 95% cluster robust confidence intervals (95% CI) and weighted importance scores (WIP) are based on model averaging across the top two models, i.e. the change in small-sample corrected Akaike Information Criterion was within 2 (ΔAICc < 2). The parameters included in each model are indicated by black dots. The intercept represents the predicted scaled degrees when all predictor variables are equal to zero or their reference (baseline) for categorical variables. For each model in the top set, we also present degrees of freedom (df), Log-likelihood (logLik), AICc, the difference in AICc from the top model (delta) and the Akaike weight, which reflects the relative support for each model.

| Response variable: Chao1 |  |  |  |  |  |  |  |  |
| --- | --- | --- | --- | --- | --- | --- | --- | --- |
| Parameter | Model | | | $\beta$ | SE | 95% CI | WIP | |
|  | 1 | 2 | 3 |  |  |  |  |  |
| (Intercept) | ● | ● | ● | 1.715 | 0.125 | (1.471, 1.960) | – |  |
| Sampling State (QLD) | ● | ● | ● | -0.763 | 0.183 | (-1.122, -0.403) | 1.00 |  |
| Sex (Male) |  | ● |  | -0.185 | 0.155 | (-0.488, 0.118) | 0.32 |  |
| Migratory status (MR) |  |  | ● | 0.105 | 0.158 | (-0.205, 0.416) | 0.20 |  |
| df | 3.00 | 4.00 | 4.00 |  |  |  |  |  |
| logLik | -167.68 | -166.94 | -167.45 |  |  |  |  |  |
| AICc | 341.71 | 342.49 | 343.49 |  |  |  |  |  |
| delta | 0.00 | 0.78 | 1.79 |  |  |  |  |  |
| weight | 0.48 | 0.32 | 0.20 |  |  |  |  |  |
| Response variable: Richness |  |  |  |  |  |  |  |  |
| Parameter | Model | | | | $\beta$ | SE | 95% CI | WIP |
|  | 1 | 2 | 3 | 4 |  |  |  |  |
| (Intercept) | ● | ● | ● | ● | 5.333 | 0.630 | (4.097, 6.568) | – |
| Sampling State (QLD) | ● | ● | ● | ● | -3.165 | 0.732 | (-4.599, -1.730) | 1.00 |
| Migratory status (MR) |  | ● |  | ● | 0.765 | 0.711 | (-0.629, 2.159) | 0.38 |
| Sex (Male) |  |  | ● | ● | -0.762 | 0.715 | (-2.163, 0.639) | 0.37 |
| df | 3.00 | 4.00 | 4.00 | 5.00 |  |  |  |  |
| logLik | -193.16 | -192.60 | -192.61 | -191.89 |  |  |  |  |
| AICc | 392.64 | 393.75 | 393.77 | 394.62 |  |  |  |  |
| delta | 0.00 | 0.10 | 1.12 | 1.97 |  |  |  |  |
| weight | 0.40 | 0.23 | 0.23 | 0.15 |  |  |  |  |

**Fig. 3.**
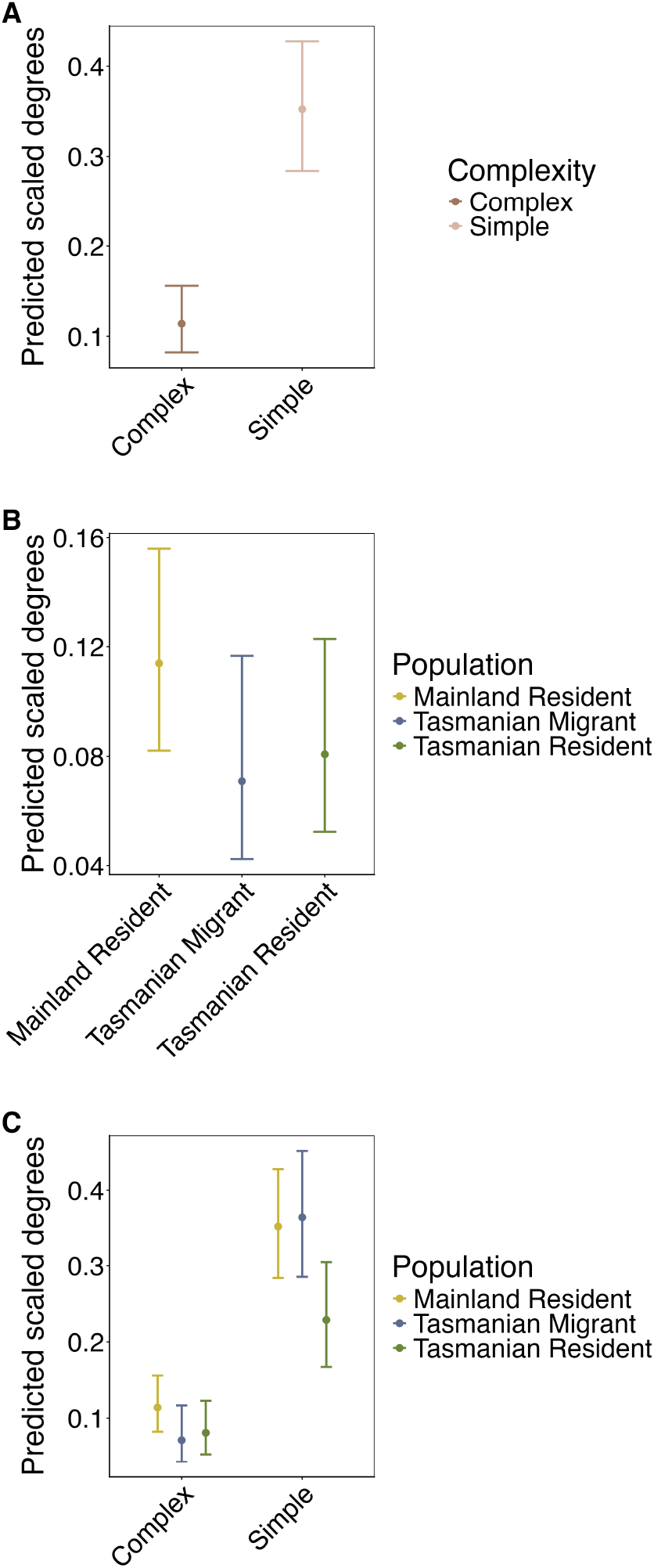
Marginal predicted effects of A) Chao1, B) Observed richness for Tasmanian migrants and mainland residents sampled in New South Wales (NSW) and Queensland (QLD). Predictions are based on an average of the top models within <2 AICc.

We found four models within <2 AICc when testing whether parasite life cycle had an impact on richness of parasites infecting the three host populations (Table S6). Life cycle complexity and population were included in all five of the top models, with both explanatory variables having a weight importance score of 1. Simple life cycle parasites infected more individuals than complex life cycle parasites (Figure. 4A). The mainland residents carried the most parasite infections, and Tasmanian residents and migrants carried the least (Figure. 4B), although there was some uncertainty in estimating this effect (Table 2). The interaction term between population and life cycle complexity was also included in three of the four top models and had a weighted importance score of 0.82. The number of simple life cycle parasites infecting Tasmanian residents was substantially lower than migrants or mainland residents (Figure. 4C). Sampling date and sex were included in one of the top models for each, and were attributed WIPs of 0.19 and 0.18, respectively. However, the 95% CI indicated some uncertainty in ascertaining the size and direction of their effect on parasite infections. We did not find substantial evidence that body condition impacted the parasite infections detected in individual birds (Table S9). The proportional similarity analysis indicated that three parasites (*Lankestrella, Plasmodium* and *Leucocytozoon*) were less evenly shared across the network than would be expected under a fixed-fixed null model (Figure. 5). These three parasites have complex life cycles. *Eimeria* was more widely distributed in the network than would be expected by chance; however, there were only two cases where *Eimeria* was detected (Figure. 1B).

**Table 2.** Model-averaged estimates for General Linear Model predicting scaled degrees (extracted from the bipartite network, representing proportion of parasite genera with simple and complex life cycles detected per host individual). Coefficient estimates (β), standard errors (SE), 95% cluster robust confidence intervals (95% CI) and weighted importance scores (WIP) are based on model averaging across the top four models, i.e. the change in small-sample corrected Akaike Information Criterion was within 2 (ΔAICc < 2). The parameters included in each model are indicated by black dots. The intercept represents the predicted scaled degrees when all predictor variables are equal to zero or their reference (baseline) for categorical variables. For each model in the top set, we also present degrees of freedom (df), Log-likelihood (logLik), AICc, the difference in AICc from the top model (delta) and the Akaike weight, which reflects the relative support for each model.

| Parameter | Model | | | | $\beta$ | SE | 95% CI | WIP |
| --- | --- | --- | --- | --- | --- | --- | --- | --- |
|  | 1 | 2 | 3 | 4 |  |  |  |  |
| (Intercept) |  | • | • | • | -2.518 | 0.343 | (-3.190, -1.846) | NA |
| Life cycle complexity (Simple) | • | • | • | • | 2.015 | 0.356 | (1.317, 2.713) | 1.00 |
| Population (Mainland Resident) | • | • | • | • | 0.520 | 0.340 | (-0.146, 1.186) | 1.00 |
| Population (Tasmanian Resident) |  | • | • | • | 0.140 | 0.396 | (-0.636, 0.916) | 1.00 |
| Life cycle complexity (Simple) × Population (Mainland Resident) | • | • |  | • | -0.700 | 0.390 | (-1.465, 0.064) | 0.82 |
| Life cycle complexity (Simple) × Population (Tasmanian Resident) | • | • |  | • | -0.977 | 0.423 | (-1.806, -0.148) | 0.82 |
| Sampling Date |  |  | • |  | -0.001 | 0.002 | (-0.006, 0.003) | 0.19 |
| Sex (Male) |  |  |  | • | -0.068 | 0.158 | (-0.378, 0.242) | 0.18 |
| df | 6.00 | 4.00 | 7.00 | 7.00 |  |  |  |  |
| logLik | -311.13 | -313.90 | -310.98 | -311.05 |  |  |  |  |
| AICc | 634.53 | 635.92 | 636.33 | 636.45 |  |  |  |  |
| delta | 0.00 | 1.39 | 1.80 | 1.92 |  |  |  |  |
| weight | 0.37 | 0.19 | 0.15 | 0.14 |  |  |  |  |

**Fig. 4.**
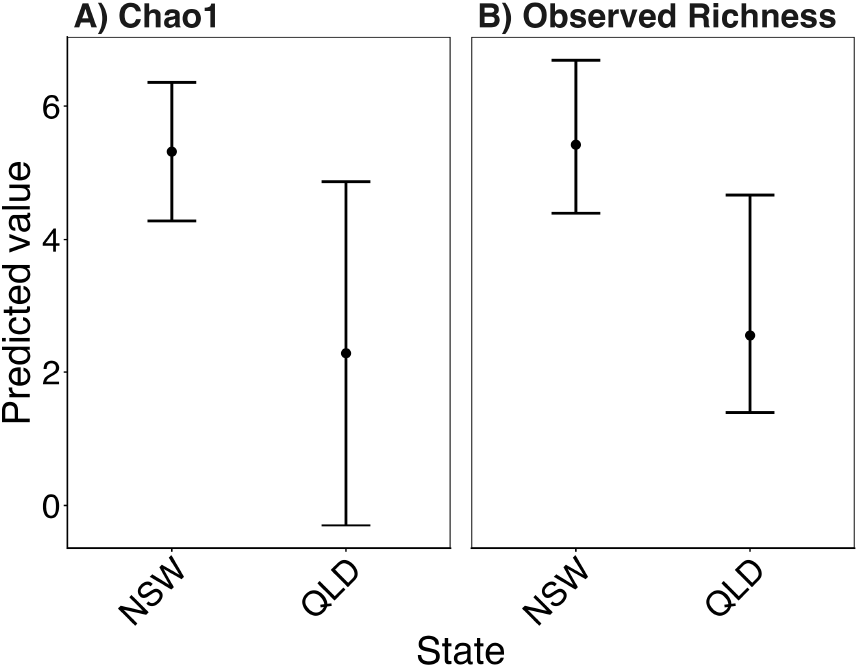
Marginal predicted effects of A) life cycle complexity, B) population and C) the interaction between life cycle complexity and population on scaled degrees extracted from the bipartite network, representing proportion of parasite genera with simple and complex life cycles detected per host individual. Predictions are based on an average of the top four models within <2 AICc.

**Fig. 5.**
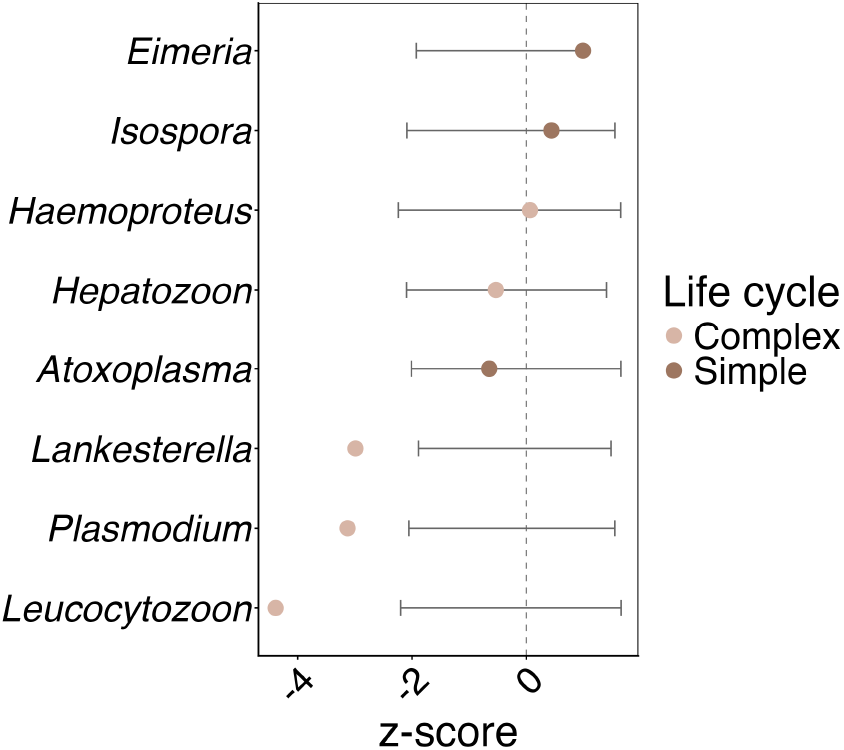
Proportional similarity z-scores for each parasite genera in the bipartite network mapping infections among 168 individuals and eight parasite genera compared to a fixed-fixed null model. Error bars indicate the range of z-scores generated in a fixed-fixed null model and the dashed line is the null expectation. Points show the observed proportional similarity values for each parasite genus and colours indicate parasite life cycle complexity.

## Discussion

By characterising a range of parasites from blood and faecal samples in a partial migrant system, we have revealed that parasite sharing facilitated by migration is influenced by life cycle complexity. While many parasite genera are shared by all three populations, migrant individuals from Tasmania appear to acquire parasite richness during the migratory period, supporting the “migratory exposure” hypothesis. This pattern is largely driven by parasites with simple life cycles which are more readily acquired by Tasmanian migrants than those with complex life cycles. Furthermore, several of the complex life cycle parasites that we detected appear to be less readily shared between the populations. Our results suggest that migrant birds may be gaining parasites from the residents they encounter on the mainland, with hosts sampled in the same mainland states displaying similar levels of parasite richness, despite differing in their migratory behaviour. Collectively, our findings from the partial migrant silvereye system expand our understanding of how transmission dynamics influence parasite sharing across the migratory range.

### Parasite sharing amongst migrants and residents is influenced by parasite life cycles

The availability of secondary hosts or vectors may constrain dispersal of parasites with complex life cycles. We detected higher parasite richness in the faeces and blood sampled from the mainland resident and Tasmanian migrant populations compared to the Tasmanian resident population. Our network analysis confirmed that these richness patterns are driven by the increase in infections of parasites with simple life cycles (particularly *Atoxoplasma* and *Isospora*) which are notably elevated in mainland resident and Tasmanian migrant populations. This result implies that complex life cycle parasites are less readily shared between hosts during the migratory cycle, a pattern observed previously and often attributed to vector availability (101; 27; 28; 29).

In the present study, *Leucocytozoon* spp. were less frequently shared among silvereye hosts than would be expected by chance. Notably, fewer infections were detected in mainland resident than Tasmanian individuals. Previous work has used *Leucocytozoon* infections to identify immigrant birds for population monitoring (102). This is possible because *Leucocytozoon* is transmitted by black flies (Diptera: Simuliidae) (103; 104), which rely on fast flowing water for larval development (105; 106) and have a relatively limited dispersal distance of up to 10 km (107). As a result, local-scale variation in vector availability impacts parasite distributions. The differing habitats of sample collection sites in Tasmania and mainland Australia may thus differentially support *Leucocytozoon* transmission. Assemblages of helminth parasites have similarly been used to track host populations, as their transmission depends on the availability of intermediate hosts such as molluscs or arthropods (108). These findings highlight that migration is more likely to facilitate the spread of simple life cycle parasites, while distributions of parasites with complex life cycles may be constrained by local ecological conditions that govern the presence and abundance of their intermediate or final hosts (109). Nonetheless, it is evident that several of the complex life cycle parasites detected in our study were broadly shared amongst the three host populations (e.g. *Haemoproteus* and *Hepatozoon*).

Differences in parasite richness became more pronounced when parasites identified in the blood and faecal samples for each host were combined, although differences among the three populations were not evident when comparing blood samples alone. The elevated richness detected in faecal samples may reflect differences in parasite life history strategies. Vector-borne parasites, which are transmitted by bites, spend part of their life cycle in host blood cells, potentially making them more readily detected in blood (110). In contrast, gut-associated parasites are more likely to rely on environmental transmission, such as excretion and ingestion of transmission stages via faecal-oral routes (111). Such environmental transmission could better facilitate parasite dispersal during migration (112; 113). Our results show that *Atoxoplasma* and *Isospora*, two parasites characterised by faecal-oral transmission are widely shared between the populations. However, it should be noted that some parasites were detected in both the faeces and blood. This was surprising as their life cycles imply restriction to a specific body compartment. The detection of parasites in host body compartments that are not associated with the parasite life cycle could reflect limitations in the availability of reference sequences (114; 53). Alternatively, it may add to the evidence that gut-associated DNA from microbes or food can be detected in the blood (115; 116; 117).

### Migrant birds gain parasites

Previous efforts to infer the directionality of parasite sharing resulting from migratory host movements have relied on multi-species comparisons of migratory and resident hosts at sites where they co-occur (27; 26; 25). Such analyses are often confounded by interspecific differences in evolutionary and ecological histories (30) and limited by sampling single points in migratory cycles. These complications may underly contradictory patterns of parasite richness and sharing between migratory and resident hosts (24; 118; 25). In our single host species study, Tasmanian migrants caught on the mainland harboured a substantially richer parasite community as measured by Chao1 and observed richness than Tasmanian resident silvereyes. The ability to compare Tasmanian migrants to their resident ‘counterfactual’ population allows us to infer that Tasmanian migrants gained parasites from mainland residents, as the latter supported richer parasite communities than Tasmanian residents. Taken together, our results provide evidence in support of the “migratory exposure” hypothesis (3) but could be further bolstered by staged sampling along the migratory route of both resident and migrating birds.

The “migratory exposure” hypothesis would further predict that parasite richness should increase with migration distance due to prolonged opportunity for exposure (119; 120). However, we observed a higher overall parasite richness in Tasmanian migrants sampled in NSW compared to QLD, despite the shorter distance travelled by migrants. It is possible that these results are explained by “migratory culling”, i.e. the death of birds that carry heavy parasite loads (18; 19), resulting in the sampled population in QLD having lower parasite richness. However, parasite richness and load for mainland resident and Tasmanian migrants were comparable in both states. This suggests that the pattern is not due to the loss of heavily infected individuals from the population. Previous work has suggested that host site use, rather than migratory distance, explained changes in helminth parasite richness in Charadriiform birds. We may similarly infer that parasite communities are determined by ecological factors operating at the local scale, causing Tasmanian migrants and mainland residents which are sampled in the same state to be more similar in their parasite richness than individuals belonging to the same population but sampled in different states.

There is a dearth of research exploring the rate at which host individuals gain and lose parasites (121; 122). It is therefore notable that, despite Tasmanian residents and migrants belonging to the same sub-species and co-occurring throughout most of the year, parasite communities were more similar between Tasmanian migrants and mainland residents. In addition, Tasmanian residents harboured the most distinct parasite communities. This is evidenced by the similarly elevated parasite richness in Tasmanian migrant and mainland residents compared to Tasmanian residents. Our LCBD analysis also showed that Tasmanian residents were characterised by a more distinctive parasite assemblage while the parasite assemblages were largely indistinguishable between migrant and mainland resident host populations. These results suggest that parasites are gained and lost by migratory birds over short timescales. Parasite life histories may further influence the rate of parasite gain and loss at the host level (122). For example, complex life cycle parasites could require prolonged periods of incubation, limiting the opportunity for transmission. Future research should consider whether transmission of parasites with complex life cycles is constrained by the length of time that migrants and residents co-occur.

## Conclusion

There are many factors that may influence the likelihood that parasites are shared as a consequence of host migration. Parasite traits can play an important role in facilitating successful sharing. These include parasite specificity (123; 124), life cycle complexity (as shown here), and life cycle flexibility, such as the ability to use paratenic hosts or to prolong stages of the life cycle (125). Virulence could reduce host survival or migratory success, thereby limiting dispersal (126; 4). Furthermore, the time required to clear infections or mount an immune response could reduce the opportunity for parasite sharing among hosts (and may also prevent successful detection) (127; 128). Parasite dispersal may thus be simultaneously influenced by a suite of factors whose varying influences should be considered when interpreting differences in parasite communities across migratory systems.

Our empirical evidence supports the “migratory exposure” hypothesis, whereby migrant birds are exposed to diverse parasite faunas during migration, resulting in a greater parasite richness in migratory birds compared to their resident counterparts. It appears that these patterns are mediated by life cycle complexity as infections with simple life cycles are particularly common in Tasmanian migrants compared to Tasmanian residents. The tendency for parasites with complex life cycles to be less readily shared throughout the network may reflect availability of their vectors or secondary hosts. Our evidence also suggests that parasite communities are determined on a local scale, with populations sampled in the same state having more similar parasite communities than those of the two Tasmanian populations. Collectively our results suggest that migration-mediated parasite sharing is constrained by the ecology and life history of the parasite. More broadly, our results emphasise the value of partial migrant systems for disentangling the ecological and evolutionary drivers of parasite transmission, and they underscore the importance of integrating host movement, parasite life history, and environmental context in future predictions of parasite spread.

## Acknowledgments

We are grateful to the many people who assisted us in identifying field sites and facilitating sample collection. In particular, we would like to thank Will Goulding, Sylvia Alexander, Ian Yeo, Brendan Doohan and many members of the Queensland Bird Research and Banding Group (Queensland); Susanne Callaghan, Alan Stuart, Rob Kyte and the National Parks and Wildlife Services (NPWS) Hunter Central Coast Branch (New South Wales); and Karina Sorrell, Menna Jones and Kat Stuart (Tasmania). We would also like to thank Vaidas Palinauskas for providing positive haemosporidian DNA extracts for use in the multiplex PCR. Thanks also to Lucy Knowles, Ewan Harney and the staff at the NERC Environmental Omics Facility (NEOF) for providing technical support for laboratory and bioinformatic work.

## Animal Studies

Ethical approval was issued to SMC (Griffith University ENV/06/20/AEC) and CY (University of Tasmania 27197). We exported our samples from Australia under permit no. PWS2022-AU-001814 and imported samples to the UK under Authorisation No. ITIMP21.1083. We thank the National Parks and Wildlife Service for New South Wales (Scientific Licence SL102647 from Department of Planning, Industry and Environment, NSW, Australia, issued to SMC), Queensland (Research permit WA0042422 from Department of Environment and Science, Queensland, Australia issued to SMC) and Tasmania (Permit No. FA22367 from Department of Natural Resources and Environment, Tasmania, Australia and Scientific Research permit SR-2022-5283, Hobart City Council, Tasmania, Australia issued to CY) for sampling permissions. We recognise ABBBS for issuing a project licence to SMC.

## Data Availability

Raw sequence reads will be deposited in the SRA upon acceptance. The code used to generate the results is available here: https://github.com/sarah-nichols/migratory-silvereye-parasites

## Author Contributions

Conceptualisation and research design: SN, SMC and BO. Investigation: SN, AE, CMY, JA, JC, JL, GL, and SMC conducted fieldwork to collect samples; SN conducted the lab work. Formal analysis: SN analysed and visualised the data. Supervision: SMC and BO. Writing: SN wrote the original draft with input from BO and SMC. All authors provided feedback on the manuscript during the drafting process.

