## Supplementary Material for "Life cycle complexity shapes parasite sharing amongst migratory and resident hosts"

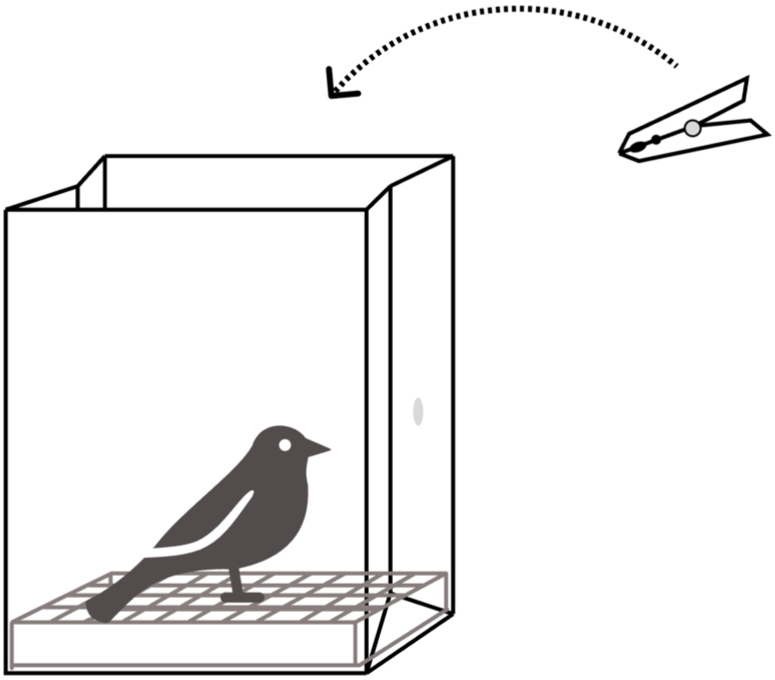

**Figure S1.** Illustration of faecal sampling protocol: flat-bottomed paper bag lined with sterile weigh-boats covered with wire mesh, ventilated by puncturing the sides of the paper bag. Pegs were used to secure the top of the bag.

**Table S1.** List of accession numbers used to build the maximum-likelihood phylogenetic tree inferred using RAxML-NG. These accession numbers were adapted from Rapp and Wolf (2024).

| Accession | Genera |
|---|---|
| AF029763.1 | *Balantidium* |
| AF043355.1 | *Bangia* |
| AF043364.2 | *Pseudobangia* |
| L26189.1 | *Erythrotrichia* |
| AF139462.1 | *Rhodochaete* |
| U25637.1 | *Ichthyophonus* |
| L29455.1 | *Choanoflagellate* |
| AF394528.1 | *Microsporidium* |
| U33180.1 | *Psorospermium* |
| D63696.1 | *Aspergillus* |
| X04971.1 | *Neurospora* |
| X54864.1 | *Pododpora* |
| U07367.1 | *Plasmodium* |
| X12708.1 | *Pneumocystis* |
| X15590.1 | *Fragaria* |
| GU476424.1 | *Eucalyptus* |
| X74753.1 | *Genicularina* |
| KM020091.1 | *Uronema* |
| X77692.1 | *Euglypha* |
| U09048.1 | *Alexandrium* |
| X59604.1 | *Babesia* |
| EF551335.1 | *Babesia* |
| KT361081.1 | *Cytauxzoon* |
| MT904044.1 | *Cytauxzoon* |
| MT904037.1 | *Cytauxzoon* |
| XR_009718807.1 | *Theileria* |
| XR_696411.1 | *Theileria* |
| FJ717705.1 | *Babesia* |
| DQ200887.1 | *Babesia* |
| MH615006.1 | *Hepatozoon* |
| MG136687.1 | *Hepatozoon* |
| KF939628.1 | *Hepatozoon* |
| KX507249.1 | *Haemogregarina* |
| KX507246.1 | *Haemogregarina* |
| U16159.1 | *Neospora* |
| EF472967.1 | *Toxoplasma* |
| XR_001974082.1 | *Toxoplasma* |

| | |
|---|---|
| L37415.1 | *Toxoplasma* |
| XR_003828656.1 | *Besnoitia* |
| AY665399.2 | *Besnoitia* |
| GU479632.1 | *Besnoitia* |
| KU341121.1 | *Sarcocystis* |
| MH590230.1 | *Sarcocystis* |
| KU341123.1 | *Sarcocystis* |
| KU180248.1 | *Lankesterella* |
| MF167545.1 | *Lankesterella* |
| AY331569.1 | *Atoxoplasma* |
| KT224380.1 | *Isospora* |
| KY801685.1 | *Isospora* |
| AF080614.1 | *Eimeria* |
| DQ136185.1 | *Eimeria* |
| EF210323.1 | *Eimeria* |
| AF080611.1 | *Lankesterella* |
| L31922.1 | *Babesia* |
| AF129883.1 | *Ophriocystis* |
| AJ535150.1 | *Achnanthes* |
| AJ535158.1 | *Amphora* |
| AF374482.2 | *Thalassiosira* |
| AJ535152.1 | *Pseudogomphonema* |
| AJ535155.1 | *Sellaphora* |
| X77703.1 | *Rhaphoneis* |
| AJ535140.1 | *Thalassionema* |
| X74752.1 | *Staurastrum* |
| AY216906.1 | *Grammatophora* |
| Y10569.1 | *Lithodesmium* |
| X85396.1 | *Thalassiosira* |
| X85395.1 | *Skeletonema* |
| AF374481.2 | *Thalassiosira* |
| AF525667.1 | *Lampriscus* |
| AJ535188.1 | *Pleurosira* |
| AF123595.1 | *Bolidomonas* |
| X85404.2 | *Aulacoseira* |
| AJ535186.1 | *Aulacoseira* |
| AJ535175.1 | *Leptocylindrus* |
| L27634.1 | *Labyrinthuloides* |
| X85399.2 | *Lauderia* |
| M54939.1 | *Lagenidium* |
| X85402.2 | *Melosira* |
| AF164133.1 | *Gastrostyla* |
| AF164128.1 | *Pleurotricha* |

| | |
|---|---|
| AF164127.1 | *Paraurostyla* |
| AF164135.1 | *Cyrtohymena* |
| X53486.1 | *Oxytricha* |
| AF164122.1 | *Oxytricha* |
| U23936.1 | *Allomyces* |
| Z14142.1 | *Palmaria* |
| AF132294.1 | *Balbiania* |
| AF026040.1 | *Audouinella* |
| AF342743.1 | *Nemalionopsis* |
| L26190.1 | *Gelidium* |
| U33133.1 | *Halymenia* |
| U09621.1 | *Rhodymenia* |
| U09618.1 | *Plocamiocolax* |
| M33639.1 | *Gracilariopsis* |
| L26183.1 | *Ceramium* |
| Z14139.1 | *Ahnfeltia* |
| AF006089.1 | *Rhodogorgon* |
| Z36893.1 | *Crossodonthina* |
| K03432.1 | *Homo* |
| NR_046237.3 | *Rattus* |
| XR_010350767.1 | *Passer* |
| L23799.1 | *Nucleospora* |
| KJ736741.1 | *Gregarina* |
| LR814087.1 | *Gregarina* |
| AY117418.1 | *Jakoba* |
| AY117417.1 | *Reclinomonas* |
| KU664396.1 | *Gregarina* |
| M19172.1 | *Plasmodium* |
| MK650492.1 | *Plasmodium* |
| MK650552.1 | *Plasmodium* |
| U07367.1 | *Plasmodium* |
| MK650858.1 | *Leucocytozoon* |

**Table S2.** Parasites identified in 168 silvereye individuals comprising 53 Tasmanian residents, 56 Tasmanian migrants and 59 mainland residents with information about their previously recorded hosts and life cycle complexity.

| Genera | Host | Life cycle | Source |
|---|---|---|---|
| *Eimeria* | Vertebrates | Simple | López-Osorio *et al.* (2020) |
| *Isospora* | Vertebrates | Simple | Fayer (1980); Box (1981); Parsa *et al.* (2023) |
| *Atoxoplasma* | Vertebrates | Simple | Levine (1982) |
| *Haemoproteus* | Vertebrates | Complex | Valkiunas (2004) |
| *Plasmodium* | Vertebrates | Complex | Valkiunas (2004) |
| *Leucocytozoon* | Vertebrates | Complex | Valkiunas (2004) |
| *Lankesterella* | Vertebrates | Complex | Keckeisen *et al.* (2024) |
| *Hepatozoon* | Vertebrates | Complex | Bennett *et al.* (1992) |

**Table S3.** Pairwise Variance inflation factors (VIFs) calculated for each predictor variable considered for inclusion in the GLMs.

| Term | VIF |
|---|---|
| Model: GLM testing sampling state impact on parasite community of residents and Tasmanian migrants | |
| Population | 1.06 |
| Sex | 1.06 |
| Sampling Date | 4.28 |
| State | 4.16 |
| Model: GLM testing life cycle complexity impact on parasite sharing | |
| Population | 1.06 |
| Life Cycle Complexity | 1.00 |
| Sex | 1.00 |
| Sampling Date | 1.06 |
| Model: GLM testing body condition impact on parasite community | |
| Body condition | 1.35 |
| Population | 1.47 |
| Sex | 1.04 |
| Sampling Date | 1.13 |

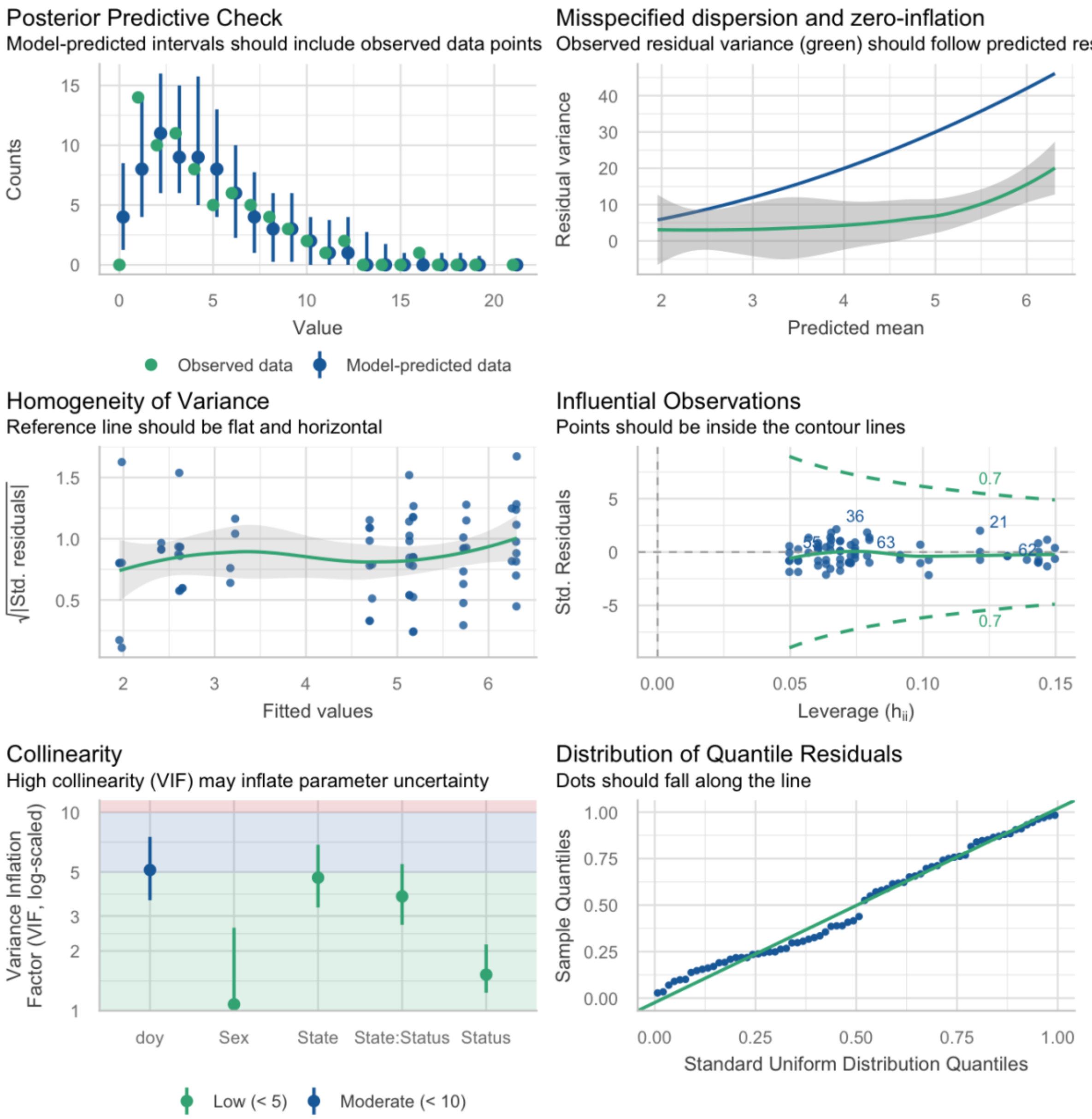


**Figure S2.** Model fit diagnostic panel produced using the performance package (Lüdecke *et al.*, 2021) testing Chao1 richness according to population, state, sex and date of sampling in Tasmanian migrants and mainland residents

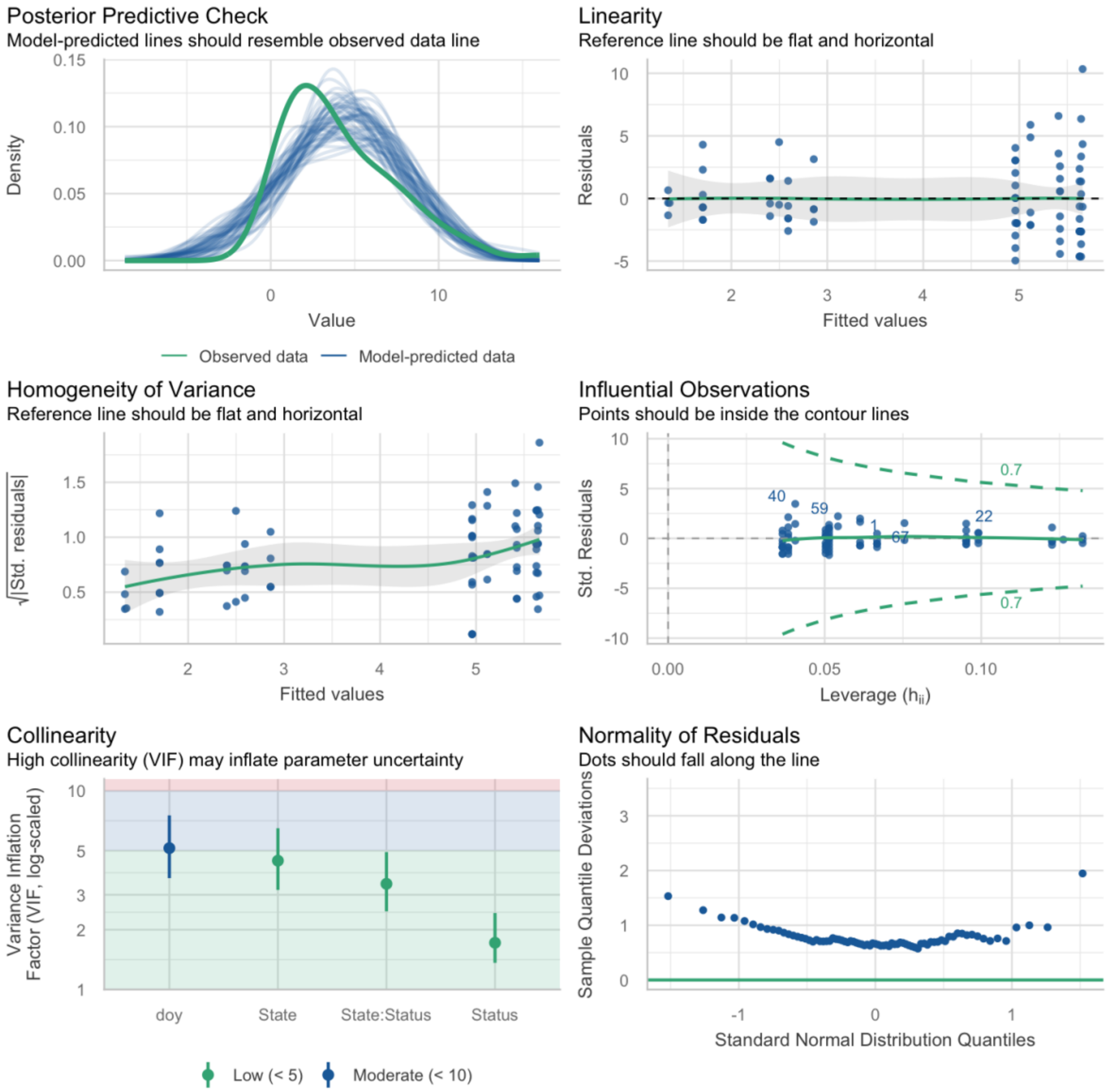


**Figure S3.** Model fit diagnostic panel produced using the performance package (Lüdecke *et al.*, 2021) testing observed richness according to population, state, sex and date of sampling in Tasmanian migrants and mainland residents.

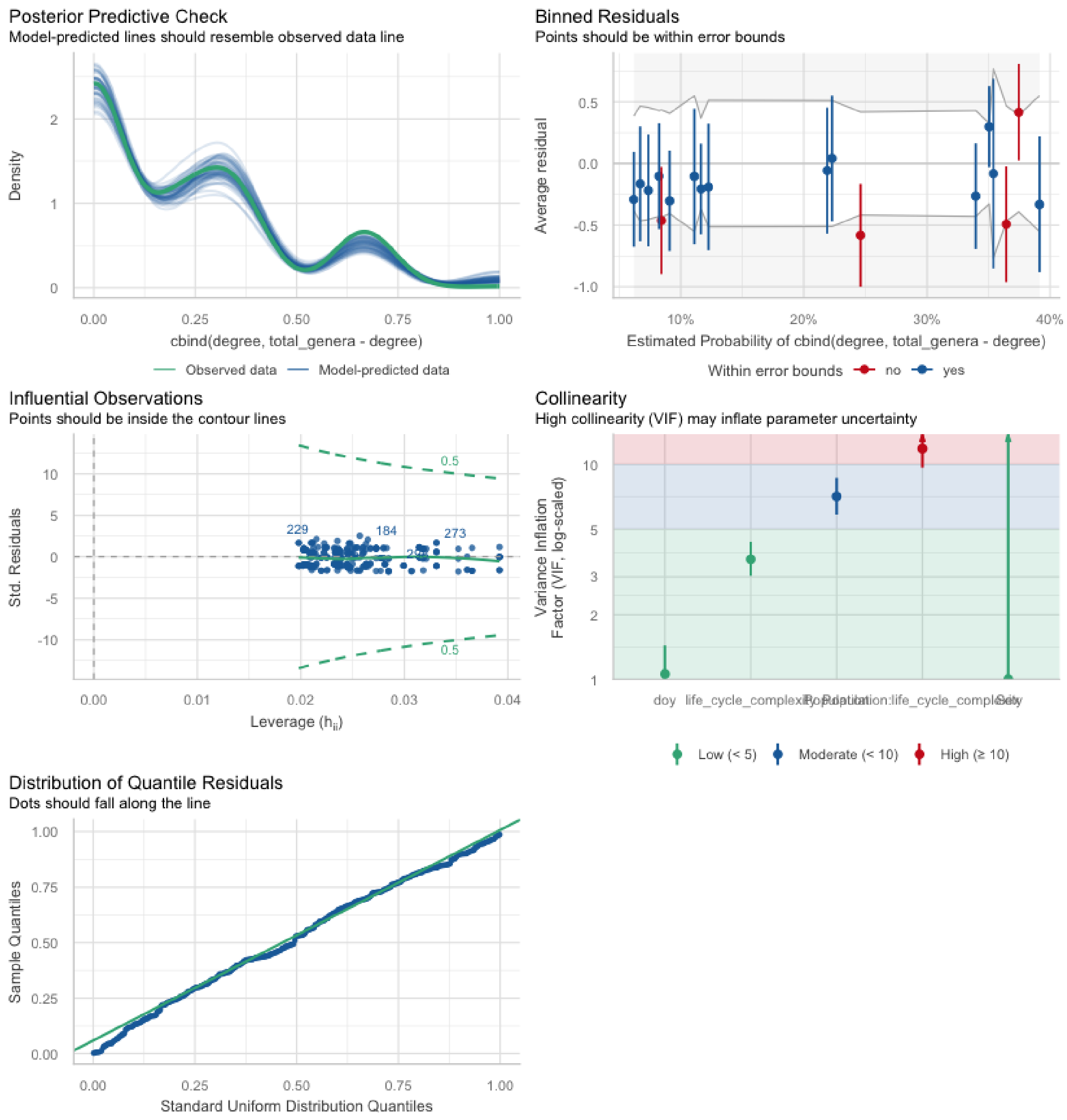


**Figure S4.** Model fit diagnostic panel produced using the performance package (Lüdecke *et al.*, 2021) for GLM predicting scaled degrees extracted from a bipartite network (i.e. the number of links to each parasite genera with simple and complex life cycles per host individual, standardised for total parasite genera).

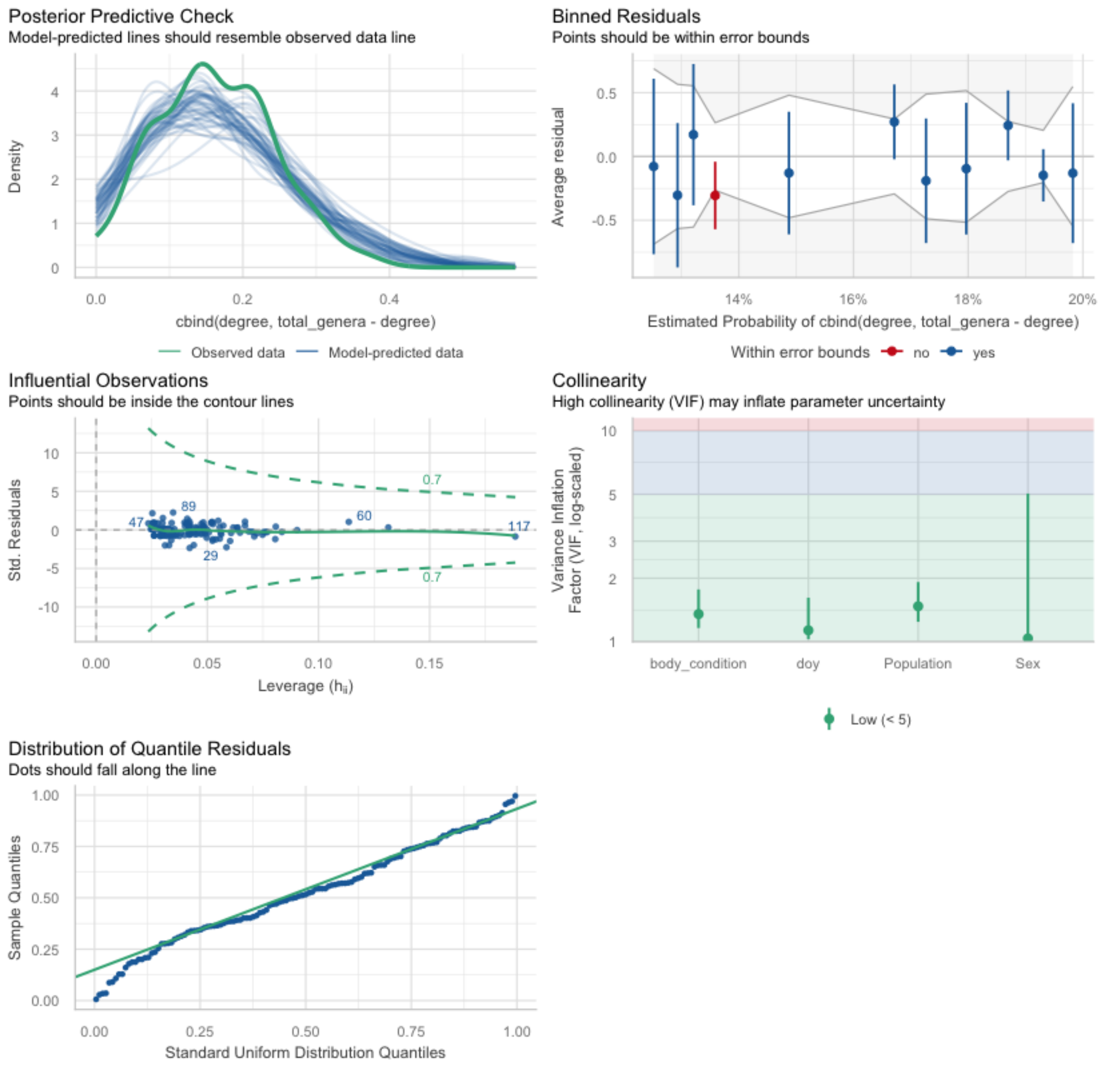


**Figure S5**. Model fit diagnostic panel produced using the performance package (Lüdecke *et al.*, 2021) for GLM with a binomial error distribution predicting scaled degrees (extracted from a bipartite network, representing parasite infections standardised according to total parasites detected) according to body condition, population, sex and date of sampling.

**Table S4.** Model selection results for GLM predicting Chao1 richness per hosts for mainland residents and Tasmanian migrants sampled in NSW and QLD. Each row represents an individual model and includes parameter estimates (intercept and coefficients), degrees of freedom (df), log-likelihood (logLik), Akaike Information Criterion corrected for small sample size (AICc), delta AICc (ΔAICc), and Akaike weights. NA values indicate that the parameter was not included in the model.

| | (Intercept) | Date of Sampling | Sex | State | Status | State x Status | df | logLik | AICc | delta | weight |
|---|---|---|---|---|---|---|---|---|---|---|---|
| 5 | 1.70 | NA | NA | + | NA | NA | 3 | -167.68 | 341.71 | 0.00 | 0.26 |
| 7 | 1.80 | NA | + | + | NA | NA | 4 | -166.94 | 342.49 | 0.78 | 0.18 |
| 13 | 1.63 | NA | NA | + | + | NA | 4 | -167.45 | 343.49 | 1.79 | 0.11 |
| 6 | 2.00 | 0.00 | NA | + | NA | NA | 4 | -167.65 | 343.90 | 2.19 | 0.09 |
| 15 | 1.72 | NA | + | + | + | NA | 5 | -166.57 | 344.04 | 2.33 | 0.08 |
| 8 | 1.79 | 0.00 | + | + | NA | NA | 5 | -166.94 | 344.80 | 3.09 | 0.06 |
| 29 | 1.66 | NA | NA | + | + | + | 5 | -167.27 | 345.46 | 3.75 | 0.04 |
| 2 | 4.17 | -0.02 | NA | NA | NA | NA | 3 | -169.55 | 345.46 | 3.75 | 0.04 |
| 14 | 1.81 | 0.00 | NA | + | + | NA | 5 | -167.44 | 345.79 | 4.08 | 0.03 |
| 31 | 1.75 | NA | + | + | + | + | 6 | -166.43 | 346.16 | 4.45 | 0.03 |
| 16 | 1.50 | 0.00 | + | + | + | NA | 6 | -166.55 | 346.40 | 4.69 | 0.03 |
| 4 | 4.12 | -0.02 | + | NA | NA | NA | 4 | -169.13 | 346.85 | 5.15 | 0.02 |
| 10 | 4.11 | -0.02 | NA | NA | + | NA | 4 | -169.48 | 347.55 | 5.84 | 0.01 |
| 30 | 2.25 | 0.00 | NA | + | + | + | 6 | -167.19 | 347.68 | 5.97 | 0.01 |
| 32 | 1.83 | 0.00 | + | + | + | + | 7 | -166.43 | 348.62 | 6.91 | 0.01 |
| 12 | 4.04 | -0.02 | + | NA | + | NA | 5 | -168.97 | 348.86 | 7.15 | 0.01 |
| 3 | 1.66 | NA | + | NA | NA | NA | 3 | -175.10 | 356.55 | 14.84 | 0.00 |
| 1 | 1.51 | NA | NA | NA | NA | NA | 2 | -176.30 | 356.78 | 15.07 | 0.00 |
| 11 | 1.58 | NA | + | NA | + | NA | 4 | -174.77 | 358.13 | 16.42 | 0.00 |
| 9 | 1.45 | NA | NA | NA | + | NA | 3 | -176.14 | 358.64 | 16.93 | 0.00 |

**Table S5**. Model selection results for GLM predicting observed richness per hosts for mainland residents and Tasmanian migrants sampled in NSW and QLD. Each row represents an individual model and includes parameter estimates (intercept and coefficients), degrees of freedom (df), log-likelihood (logLik), Akaike Information Criterion corrected for small sample size (AICc), delta AICc (ΔAICc), and Akaike weights. NA values indicate that the parameter was not included in the model.

| | (Intercept) | Date of Sampling | Sex | State | Status | State x Status | df | logLik | AICc | delta | weight |
|---|---|---|---|---|---|---|---|---|---|---|---|
| 5 | 5.34 | NA | NA | + | NA | NA | 3.00 | -193.16 | 392.64 | 0.00 | 0.24 |
| 13 | 4.90 | NA | NA | + | + | NA | 4.00 | -192.60 | 393.75 | 1.10 | 0.14 |
| 7 | 5.76 | NA | + | + | NA | NA | 4.00 | -192.61 | 393.77 | 1.12 | 0.14 |
| 15 | 5.32 | NA | + | + | + | NA | 5.00 | -191.89 | 394.62 | 1.97 | 0.09 |
| 6 | 6.78 | -0.01 | NA | + | NA | NA | 4.00 | -193.12 | 394.80 | 2.16 | 0.08 |
| 29 | 5.00 | NA | NA | + | + | + | 5.00 | -192.55 | 395.94 | 3.29 | 0.05 |
| 14 | 6.03 | -0.01 | NA | + | + | NA | 5.00 | -192.57 | 395.99 | 3.35 | 0.05 |
| 8 | 6.00 | 0.00 | + | + | NA | NA | 5.00 | -192.60 | 396.05 | 3.41 | 0.04 |
| 2 | 15.91 | -0.08 | NA | NA | NA | NA | 3.00 | -194.98 | 396.29 | 3.65 | 0.04 |
| 31 | 5.41 | NA | + | + | + | + | 6.00 | -191.84 | 396.89 | 4.24 | 0.03 |
| 16 | 5.02 | 0.00 | + | + | + | NA | 6.00 | -191.88 | 396.97 | 4.32 | 0.03 |
| 10 | 15.33 | -0.08 | NA | NA | + | NA | 4.00 | -194.51 | 397.58 | 4.94 | 0.02 |
| 4 | 15.91 | -0.07 | + | NA | NA | NA | 4.00 | -194.76 | 398.07 | 5.43 | 0.02 |
| 30 | 7.28 | -0.02 | NA | + | + | + | 6.00 | -192.48 | 398.15 | 5.51 | 0.02 |
| 12 | 15.27 | -0.07 | + | NA | + | NA | 5.00 | -194.19 | 399.22 | 6.58 | 0.01 |
| 32 | 5.89 | 0.00 | + | + | + | + | 7.00 | -191.84 | 399.30 | 6.66 | 0.01 |
| 1 | 4.22 | NA | NA | NA | NA | NA | 2.00 | -202.13 | 408.42 | 15.78 | 0.00 |
| 3 | 4.74 | NA | + | NA | NA | NA | 3.00 | -201.49 | 409.32 | 16.67 | 0.00 |
| 9 | 3.72 | NA | NA | NA | + | NA | 3.00 | -201.51 | 409.34 | 16.70 | 0.00 |
| 11 | 4.24 | NA | + | NA | + | NA | 4.00 | -200.69 | 409.94 | 17.30 | 0.00 |

**Table S6.** Model selection results for GLM predicting scaled degrees from simple and complex life cycle parasites in a bipartite network. Each row represents an individual model and includes parameter estimates (intercept and coefficients), degrees of freedom (df), log-likelihood (logLik), Akaike Information Criterion corrected for small sample size (AICc), delta AICc (ΔAICc), and Akaike weights. NA values indicate that the parameter was not included in the model.

| | (Intercept) | Sampling Date | Life cycle complexity | Population | Sex | Life cycle complexity ×Population | df | logLik | AICc | delta | weight |
|---|---|---|---|---|---|---|---|---|---|---|---|
| 23 | -2.64 | NA | + | + | NA | + | 6 | -308.59 | 629.44 | 0.00 | 0.35 |
| 24 | -2.39 | 0.00 | + | + | NA | + | 7 | -308.43 | 631.22 | 1.77 | 0.14 |
| 7 | -2.27 | NA | + | + | NA | NA | 4 | -311.57 | 631.27 | 1.82 | 0.14 |
| 31 | -2.60 | NA | + | + | + | + | 7 | -308.50 | 631.35 | 1.91 | 0.13 |
| 8 | -2.03 | 0.00 | + | + | NA | NA | 5 | -311.42 | 633.02 | 3.58 | 0.06 |
| 32 | -2.35 | 0.00 | + | + | + | + | 8 | -308.34 | 633.14 | 3.70 | 0.05 |
| 15 | -2.23 | NA | + | + | + | NA | 5 | -311.48 | 633.15 | 3.70 | 0.05 |
| 3 | -2.31 | NA | + | NA | NA | NA | 2 | -315.43 | 634.89 | 5.45 | 0.02 |
| 16 | -1.99 | 0.00 | + | + | + | NA | 6 | -311.33 | 634.92 | 5.48 | 0.02 |
| 11 | -2.26 | NA | + | NA | + | NA | 3 | -315.28 | 636.64 | 7.20 | 0.01 |
| 4 | -2.20 | 0.00 | + | NA | NA | NA | 3 | -315.39 | 636.85 | 7.41 | 0.01 |
| 12 | -2.16 | 0.00 | + | NA | + | NA | 4 | -315.25 | 638.63 | 9.19 | 0.00 |
| 5 | -1.51 | NA | NA | + | NA | NA | 3 | -363.75 | 733.57 | 104.13 | 0.00 |
| 6 | -1.28 | 0.00 | NA | + | NA | NA | 4 | -363.61 | 735.34 | 105.89 | 0.00 |
| 13 | -1.47 | NA | NA | + | + | NA | 4 | -363.66 | 735.45 | 106.01 | 0.00 |
| 1 | -1.55 | NA | NA | NA | NA | NA | 1 | -367.29 | 736.59 | 107.14 | 0.00 |
| 14 | -1.25 | 0.00 | NA | + | + | NA | 5 | -363.53 | 737.24 | 107.80 | 0.00 |
| 9 | -1.51 | NA | NA | NA | + | NA | 2 | -367.16 | 738.35 | 108.91 | 0.00 |
| 2 | -1.45 | 0.00 | NA | NA | NA | NA | 2 | -367.25 | 738.54 | 109.10 | 0.00 |
| 10 | -1.41 | 0.00 | NA | NA | + | NA | 3 | -367.13 | 740.33 | 110.89 | 0.00 |
| 23 | -2.64 | NA | + | + | NA | + | 6 | -308.59 | 629.44 | 0.00 | 0.35 |
| 24 | -2.39 | 0.00 | + | + | NA | + | 7 | -308.43 | 631.22 | 1.77 | 0.14 |
| 7 | -2.27 | NA | + | + | NA | NA | 4 | -311.57 | 631.27 | 1.82 | 0.14 |
| 31 | -2.60 | NA | + | + | + | + | 7 | -308.50 | 631.35 | 1.91 | 0.13 |
| 8 | -2.03 | 0.00 | + | + | NA | NA | 5 | -311.42 | 633.02 | 3.58 | 0.06 |
| 32 | -2.35 | 0.00 | + | + | + | + | 8 | -308.34 | 633.14 | 3.70 | 0.05 |
| 15 | -2.23 | NA | + | + | + | NA | 5 | -311.48 | 633.15 | 3.70 | 0.05 |
| 3 | -2.31 | NA | + | NA | NA | NA | 2 | -315.43 | 634.89 | 5.45 | 0.02 |
| 16 | -1.99 | 0.00 | + | + | + | NA | 6 | -311.33 | 634.92 | 5.48 | 0.02 |
| 11 | -2.26 | NA | + | NA | + | NA | 3 | -315.28 | 636.64 | 7.20 | 0.01 |
| 4 | -2.20 | 0.00 | + | NA | NA | NA | 3 | -315.39 | 636.85 | 7.41 | 0.01 |
| 12 | -2.16 | 0.00 | + | NA | + | NA | 4 | -315.25 | 638.63 | 9.19 | 0.00 |
| 5 | -1.51 | NA | NA | + | NA | NA | 3 | -363.75 | 733.57 | 104.13 | 0.00 |
| 6 | -1.28 | 0.00 | NA | + | NA | NA | 4 | -363.61 | 735.34 | 105.89 | 0.00 |
| 13 | -1.47 | NA | NA | + | + | NA | 4 | -363.66 | 735.45 | 106.01 | 0.00 |

| | | | | | | | | | | | |
|---|---|---|---|---|---|---|---|---|---|---|---|
| 1 | -1.55 | NA | NA | NA | NA | NA | 1 | -367.29 | 736.59 | 107.14 | 0.00 |
| 14 | -1.25 | 0.00 | NA | + | + | NA | 5 | -363.53 | 737.24 | 107.80 | 0.00 |
| 9 | -1.51 | NA | NA | NA | + | NA | 2 | -367.16 | 738.35 | 108.91 | 0.00 |
| 2 | -1.45 | 0.00 | NA | NA | NA | NA | 2 | -367.25 | 738.54 | 109.10 | 0.00 |
| 10 | -1.41 | 0.00 | NA | NA | + | NA | 3 | -367.13 | 740.33 | 110.89 | 0.00 |

**Table S7.** Model selection results for GLM predicting scaled degrees extracted from a bipartite network (i.e. the number of links to each parasite genera per host individual, standardised for total parasite genera). Each row represents an individual model and includes parameter estimates (intercept and coefficients), degrees of freedom (df), log-likelihood (logLik), Akaike Information Criterion corrected for small sample size (AICc), delta AICc (ΔAICc), and Akaike weights. NA values indicate that the parameter was not included in the model.

| | (Intercept) | Body Condition | Sampling Date | Population | Sex | df | logLik | AICc | delta | weight |
|---|---|---|---|---|---|---|---|---|---|---|
| 5 | -1.46 | NA | NA | + | NA | 3 | -196.35 | 398.89 | 0.00 | 0.27 |
| 6 | -1.49 | -0.06 | NA | + | NA | 4 | -196.13 | 400.58 | 1.69 | 0.11 |
| 7 | -1.09 | NA | 0.00 | + | NA | 4 | -196.19 | 400.69 | 1.81 | 0.11 |
| 13 | -1.47 | NA | NA | + | + | 4 | -196.34 | 401.00 | 2.12 | 0.09 |
| 2 | -1.65 | -0.15 | NA | NA | NA | 2 | -198.56 | 401.21 | 2.33 | 0.08 |
| 4 | -0.90 | -0.14 | 0.00 | NA | NA | 3 | -197.88 | 401.95 | 3.07 | 0.06 |
| 8 | -1.13 | -0.06 | 0.00 | + | NA | 5 | -195.98 | 402.44 | 3.56 | 0.05 |
| 14 | -1.50 | -0.06 | NA | + | + | 5 | -196.11 | 402.71 | 3.82 | 0.04 |
| 15 | -1.09 | NA | 0.00 | + | + | 5 | -196.17 | 402.83 | 3.94 | 0.04 |
| 3 | -0.67 | NA | -0.01 | NA | NA | 2 | -199.48 | 403.06 | 4.18 | 0.03 |
| 10 | -1.68 | -0.16 | NA | NA | + | 3 | -198.50 | 403.20 | 4.32 | 0.03 |
| 1 | -1.65 | NA | NA | NA | NA | 1 | -200.62 | 403.26 | 4.38 | 0.03 |
| 12 | -0.91 | -0.14 | 0.00 | NA | + | 4 | -197.79 | 403.89 | 5.01 | 0.02 |
| 16 | -1.14 | -0.06 | 0.00 | + | + | 6 | -195.94 | 404.57 | 5.69 | 0.02 |
| 11 | -0.67 | NA | -0.01 | NA | + | 3 | -199.44 | 405.07 | 6.19 | 0.01 |
| 9 | -1.66 | NA | NA | NA | + | 2 | -200.61 | 405.31 | 6.42 | 0.01 |

**Table S8.** Pairwise Dunn's tests for each sample type and metric calculated using ASV sequences extracted from faecal and blood samples of 160 silvereye, group 1 and 2 indicate pairwise comparison. LCBD = Local Contribution to Beta Diversity. P values adjusted with Holm correction for multiple comparisons.

| Sample Type | Metric | Group 1 | Group 2 | Statistic | *p.adj* |
|---|---|---|---|---|---|
| Blood | Chao1 | Tasmania Resident | Migrant | 0.210 | 1.000 |
| Blood | Chao1 | Tasmania Resident | Mainland Resident | -0.425 | 1.000 |
| Blood | Chao1 | Migrant | Mainland Resident | -0.629 | 1.000 |
| Blood | LCBD | Tasmania Resident | Migrant | -0.029 | 1.000 |
| Blood | LCBD | Tasmania Resident | Mainland Resident | -0.417 | 1.000 |
| Blood | LCBD | Migrant | Mainland Resident | -0.368 | 1.000 |
| Blood | Observed Richness | Tasmania Resident | Migrant | -0.817 | 0.828 |
| Blood | Observed Richness | Tasmania Resident | Mainland Resident | 0.488 | 0.828 |
| Blood | Observed Richness | Migrant | Mainland Resident | 1.341 | 0.540 |
| Combined | Chao1 | Tasmania Resident | Migrant | 2.434 | 0.045 |
| Combined | Chao1 | Tasmania Resident | Mainland Resident | 2.404 | 0.045 |
| Combined | Chao1 | Migrant | Mainland Resident | -0.112 | 0.911 |
| Combined | LCBD | Tasmania Resident | Migrant | -2.398 | 0.049 |
| Combined | LCBD | Tasmania Resident | Mainland Resident | -1.251 | 0.422 |
| Combined | LCBD | Migrant | Mainland Resident | 1.235 | 0.422 |
| Combined | Observed Richness | Tasmania Resident | Migrant | 1.713 | 0.174 |
| Combined | Observed Richness | Tasmania Resident | Mainland Resident | 2.398 | 0.050 |
| Combined | Observed Richness | Migrant | Mainland Resident | 0.673 | 0.501 |
| Faecal | Chao1 | Tasmania Resident | Migrant | 2.626 | 0.026 |
| Faecal | Chao1 | Tasmania Resident | Mainland Resident | 1.648 | 0.199 |
| Faecal | Chao1 | Migrant | Mainland Resident | -1.154 | 0.249 |
| Faecal | LCBD | Tasmania Resident | Migrant | -2.761 | 0.017 |
| Faecal | LCBD | Tasmania Resident | Mainland Resident | -1.143 | 0.253 |
| Faecal | LCBD | Migrant | Mainland Resident | 1.856 | 0.127 |
| Faecal | Observed Richness | Tasmania Resident | Migrant | 3.090 | 0.006 |
| Faecal | Observed Richness | Tasmania Resident | Mainland Resident | 3.033 | 0.006 |
| Faecal | Observed Richness | Migrant | Mainland Resident | -0.109 | 0.913 |

**Table S9**. Model-averaged estimates for GLM predicting scaled degrees (proportion of degrees linking parasite genera to host individuals extracted from bipartite network). Coefficient estimates (β), standard errors (SE), 95% cluster robust confidence intervals (95% CI) and weighted importance scores (WIP) are based on model averaging across the top six models ($\Delta AICc < 2$). The parameters included in each model are indicated by black dots. The intercept represents the predicted scaled degrees when all predictor variables are equal to zero or their reference (baseline) for categorical variables. For each model in the top set, we also present degrees of freedom (df), Log-likelihood (logLik), small-sample corrected Akaike Information Criterion (AICc), the difference in AICc from the top model (delta) and the Akaike weight, which reflects the relative support for each model.

| Parameter | 1 | 2 | 3 | β | SE | 95% CI | WIP |
|---|---|---|---|---|---|---|---|
| (Intercept) | • | • | • | -1.384 | 0.359 | (-2.087, -0.680) | NA |
| Population (Tasmanian Migrant) | • | • | • | -0.124 | 0.165 | (-0.447, 0.199) | 1.00 |
| Population (Tasmanian Resident) | • | • | • | -0.408 | 0.160 | (-0.722, -0.094) | 1.00 |
| Body Condition | | • | | -0.058 | 0.089 | (-0.232, 0.116) | 0.23 |
| Sampling Date | | | • | -0.002 | 0.004 | (-0.011, 0.006) | 0.22 |
| df | 3.00 | 4.00 | 4.00 | | | | |
| logLik | -196.35 | -196.13 | -196.19 | | | | |
| AICc | 398.89 | 400.58 | 400.69 | | | | |
| delta | 0.00 | 1.69 | 1.81 | | | | |
| weight | 0.55 | 0.23 | 0.22 | | | | |
